# Comparative analyses of phenotypic sequences using phylogenetic trees

**DOI:** 10.1101/561167

**Authors:** Daniel S. Caetano, Jeremy M. Beaulieu

## Abstract

Phenotypic sequences are a type of multivariate trait organized structurally, such as teeth distributed along the dental arch, or temporally, such as the stages of an ontogenetic series. However, unlike other multivariate traits, the elements of a phenotypic sequence are arranged along a vector, which allows for distinct evolutionary patterns between neighboring and distant positions. In fact, sequence traits share many characteristics with molecular sequences. We implement an approach to estimate rates of trait evolution that explicitly incorporates the sequence organization of traits. We apply models to study the temporal pattern evolution of cricket calling songs. We test whether songs show autocorrelation of rates (i.e., neighboring positions along a phenotypic sequence have correlated rates of evolution), or if they are best described by rate variation independent of sequence position. Our results show that models perform well when used with sequence phenotypes even under small sample sizes. We also show that silent regions of the songs evolve faster than chirp regions, which suggests that macroevolutionary changes are faster when associated with axes of variation less constrained by multiple sources of selection. Our approach is flexible and can be applied to any multivariate trait with units organized in a sequence-like structure.

## Introduction

The diversity of phenotypes observed across the tree of life can be grouped into hierarchical levels. At the simplest level of organization, univariate traits are treated as independent units. One level higher, multiple univariate traits that covary due to genetic correlation or developmental constraints group to form modules (Olson and Miller 1958, Klingenberg and Marugán-Lobón 2013, Armbruster et al. 2014, Goswami et al. 2014, Caetano and Harmon 2018). Finally, the trait itself can be composed of multiple dimensions and, unlike modules, these dimensions cannot be easily separated into univariate parts (Adams 2014, Adams and Felice 2014, also see discussion on Caetano and Harmon 2018). Geometric morphometric data and other complex traits, such as animal vocalizations, belong to the multidimensional category because the coordinates of a point in a plane (e.g., Adams 2014, Adams and Felice 2014), or a single sound note (e.g., Otte 1992, Rohrmeier 2015), are not meaningful in isolation. However, both evolutionarily correlated traits and multidimensional phenotypes are often transformed into principal axes of variation (e.g., Klingenberg and Marugán-Lobón 2013, Genevcius et al. 2017, also see discussion on Uyeda et al. 2015) or reduced to subunits (e.g., Caetano and Machado 2013, Ligon et al. 2018) to facilitate identification of interspecific relationships, statistical modeling, and visualization.

The hierarchical organization of phenotypes plays an important role in shaping biodiversity. Phenotypic integration, for instance, enables a coordinated response to selection whereas independent traits are less evolutionarily constrained and can evolve to occupy a relatively larger region of the morphospace (Goswami et al. 2014). Unfortunately, the depth of our knowledge about evolutionary patterns and processes is not the same across these levels, since the majority of comparative studies focus on univariate traits rather than any higher level of phenotypic organization. Phenotypic sequences, for instance, are a special case of multidimensional phenotype rarely present in phylogenetic studies. Sequence traits can be defined as a series of phenotypic units distributed along a vector, which is usually composed by a series of adjacent anatomical positions (e.g., Stock 2001, Smith 2003, Peel et al. 2005, Smith et al. 2013, Zhu et al. 2017) or time steps (e.g., Richardson et al. 2001, Jeffery et al. 2002, Robillard et al. 2006, Scholes III 2008, Arnold et al. 2017). Teeth expressed along the dental arch are an example of a phenotypic sequence organized structurally. Teeth are modular structures that share characteristics, including function, within the same group which can differ from other teeth groups. Importantly, the structural organization of teeth along the dental arch can facilitate evolution of position-dependent specialization, meaning that teeth from different regions of the dental arch can show varied evolutionary trajectories (see discussion in Stock 2001). Ontogeny can also be described as a sequence trait, but organized in a vector of time steps (Richardson et al. 2001, Jeffery et al. 2002). For example, heterochrony, the evolution of novel morphologies by manipulation of the relative timing of developmental events, is an important macroevolutionary mechanism which is best described as a sequence (Jeffery et al. 2002).

The term “sequence” is broadly used as a shorthand for “molecular sequence” and, indeed, molecular sequences share many characteristics with sequence traits. For instance, models of sequence evolution estimate rates of transition between units along the positions of a vector defined by the structure of the DNA (i.e., loci). Thus, many concepts originally designed to study molecular evolution, such as the relationship between neighboring loci (Yang 1993, Yang 1995), are likely applicable to phenotypic sequences, although such a strategy has been rarely attempted (but see Robillard et al. 2006). The number of phylogenetic comparative methods aimed at the study of multivariate traits has increased recently (e.g., Klingenberg and Marugán-Lobón 2013, Clavel et al. 2015, Goswami and Finarelli 2016, Adams and Collyer 2017, Caetano and Harmon 2017, Caetano and Harmon 2018, Clavel et al. 2019). However, these methods differ from molecular evolution approaches and are not adequate for sequence traits because they do not incorporate the serial organization of the states observed at the tips of the phylogeny. In fact, most multivariate comparative methods take into account the evolutionary correlation among traits, but are agnostic to any particular ordering or ranking (Clavel et al. 2015, Caetano and Harmon 2017, Caetano and Harmon 2018).

Here we use phylogenetic comparative methods adapted from models of molecular evolution in order to study rates of evolution for sequence traits (Yang 1993, Yang 1995). We study the temporal pattern in calling songs of 11 species of field crickets (*Gryllus*, Gryllidae). These songs have a strong genetic component and are an important trait for species recognition, species reinforcement, and taxonomic identification (Otte 1992, Robillard et al. 2006, Robillard and Desutter-Grandcolas 2011, Blankers et al. 2018, Gray et al. 2018). Calling songs in crickets evolve early in speciation and are often the only trait allowing recognition of recently isolated lineages (Otte 1992, Gray and Cade 2000, Mendelson and Shaw 2005, Gray et al. 2016). Calling songs differ between species in many axes of variation, including temporal patterns comprised by sound regions (chirp) alternated by interchirp silence regions (Otte 1992, Robillard et al. 2006, Robillard and Desutter-Grandcolas 2011). We estimate rates of evolution along the sequence of calling songs using their temporal patterns and ask whether, 1) sound and silence regions evolve at distinct rates, and 2) if there is signal in the data for autocorrelation of rates.

## Methods

### Overview

We implemented a set of models to estimate rates of evolution, and evaluate hypotheses of evolutionary correlation among the positions of phenotypic sequences. These models are analogous to models of molecular evolution (e.g., Tavaré 1986, Yang 1993, Yang 1995, Sullivan and Joyce 2005, Beaulieu et al. 2019), but are conditioned on a known phylogeny. Transition rates among states can be constant across sequence positions or they can vary following two distinct patterns: either independent of position or correlated among neighboring sites. All models implemented here apply continuous-time Markov chains which are widely used in comparative methods and molecular phylogenetics (reviewed by O’Meara 2012). We used this approach to analyze the temporal patterns of male calling song sequences for 11 species of *Gryllus* crickets (Robillard et al. 2006), while taking into account phylogenetic uncertainty, sequence alignment variation, and hypotheses of homology at different levels. Finally, we conducted simulations to explore the performance of the models, especially with respect to datasets of smaller sizes (i.e., few homologous “sites”).

Models of sequence evolution are almost exclusively developed for molecular data and pose many challenges for the study of *phenotypic* sequence evolution. For example, the size of the state space is known and constant among loci in molecular sequences. This means that the dimensions of the Markov transition matrix (**Q**) can be determined *a priori* and shared across sites (Felsenstein 1981, Tavaré 1986). The states of molecular sequences are also connected by clear homology statements, which allow substitutions to be jointly estimated across loci. Complex models of nucleotide substitution, such as the Generalized time-reversible model (GTR; Tavaré 1986), are able to estimate unequal transition rates between different base pairs because homolog substitutions (e.g., between base-pairs “A” and “T”) represent the same substitution event across all loci. By contrast, the realized state space of phenotypic sequences will likely vary across positions because some positions can have only a few observed states at the tips of the phylogeny, whereas others could have many more. Importantly, a homology statement connecting states on different sequence positions of a phenotype may be absent. In the case of cricket calling song data, we were able to erect a homology statement for observed states across loci based on the repeated structure of the trait, but we also acknowledge many limitations in modeling phenotypic sequences. For instance, here we assume equal transition rates among observed states and a **Q** matrix scaled to the observed state diversity at each sequence position.

Multiple sequence alignment (MSA) is the first step in any phylogenetic inference analyses using molecular data and arguably the most important one, since it defines site-wise homologies that, in most cases, will not be re-evaluated by downstream analyses (but see Redelings and Suchard 2005). Most MSA methods can be classified as either distance-based approaches (e.g., Katoh and Standley 2013) or statistical alignment models (e.g., Redelings and Suchard 2005, Suchard and Redelings 2006). The main difference is that the former provides a single alignment that minimizes the distance between sequences following a cost matrix, whereas the latter evaluates the likelihood of a model of sequence evolution given a set of parameter values and, coupled with Bayesian estimates, allow for sampling sequence alignments in proportion to their posterior probabilities. Currently, the majority of phylogenetic studies apply distance-based methods, likely because the alignment quality provided by statistical alignments (Redelings 2014) is counterbalanced by the extreme increase in the use of computational resources (Nute and Warnow 2016).

Unlike MSA used in phylogenetic inferences, the alignment of non-molecular sequences can be inferred conditioned on a tree that has been derived from independent data. Fixing topology and branch lengths reduce the dimensionality of the problem and facilitates the use of model-based alignment approaches, which otherwise perform joint estimation of the MSA, topology, and branch lengths (Redelings and Suchard 2005, Suchard and Redelings 2006). Unfortunately, the mechanism of trait evolution in phenotypic sequences is likely to be unknown, or at least known in much less detail, than nucleotide evolution and one might argue that the use of position-independent MSA approaches, such as BAli-Phy (Suchard and Redelings 2006), would be unrealistic. Although we acknowledge the limitations of the current methods, we urge the reader to consider ours as an attempt to bridge fairly disparate disciplines and, by doing so, also highlight important avenues for future improvement (see *Current challenges and moving forward*).

### Modelling rate variation for phenotypic sequences

We first implemented a model with rates varying along the phenotypic sequence independent of position. Specifically, rates are drawn from a standard Gamma distribution — that is, the parameter α defines the shape of the distribution, which is then discretized into *k* equiprobable rate categories as described by Yang (1993). In short, this model independently distributes *k* rate scaling categories applied to a global transition matrix (**Q**) shared among all sequence positions. The transition rate is then obtained by marginalizing the *k* rate categories at each sequence position. In other words, the overall likelihood reflects the combination of rate categories that improve the probability of observing the data at each sequence position (Yang 1993). Hereafter, we will refer to this model as the ‘gamma’ model. The gamma model allows for distinct regions of the phenotypic sequence to evolve at different rates and is useful to test shifts in rates of transition along the positions of a sequence while assuming that each position is allowed to evolve at independent rates (i.e., in the absence of autocorrelation).

We also suppose that evolutionary rates may not be independent across sequence positions as a result of evolutionary correlation (i.e., autocorrelation of rates along the sequence). Yang (1995) described an extension of the gamma model that allows neighboring positions to have correlated rates of evolution by adding a single extra parameter controlling the correlation of a bivariate standard Gamma distribution; this is referred to as the auto-discrete gamma model (Yang 1995). In this model, the probability of assigning rate category *k_i_* to a position of the sequence *p_n_* is conditioned on observing category *k_j_* on the neighboring position *p_n-1_*. This conditional probability is governed by a symmetric discrete-time hidden Markov model (HMM) running along the sequence and it is independent of direction (Yang 1995). As a result, the evolutionary rate for a given position will be dependent on its neighboring positions on either side. While Yang (1995) referred to this model as the auto-discrete-gamma rates model, we refer hereafter to this model as the ‘correlated’ model. We note that the correlated model can be especially interesting in cases when elements of a phenotypic sequence are more likely to share evolutionary rates if close together along the sequence, such as stages in a courtship display (e.g., Scholes III 2008, Arnold et al. 2017) or ontogenetic development (e.g., Richardson et al. 2001, Jeffery et al. 2002). One can also expect non-independent rates of evolution on genomic studies, because genes from physically proximate regions show coordinated expression (i.e., expression piggybacking - Ghanbarian and Hurst 2015) and correlated evolution (i.e., genetic hitchhiking - Barton 2000). And, unlike the gamma model (Yang 1993), the correlated model (Yang 1995) is rarely used in phylogenetics, although the role of dependency among sites has been shown to be important in molecular sequences (see Huelsenbeck and Nielsen 1999).

### Dealing with gaps

Multiple sequence alignments produce gaps representing insertions and deletions (i.e., indels) of sequence positions. In the case of molecular sequences, gaps are simply treated as missing data. This is less straightforward with phenotypic sequences, because transitions to and from gap states actually represent the rate of losses and gains of the *observed state* in a given sequence position. It is possible, however, to partition the model to estimate separate rates for transitions among observed states as well as transitions into and out of gap “states”. For this we included two transition schemes. In the simplest case, we estimate a single transition rate between all states in the data, agnostic of gap states. Alternatively, transitions between observed states and transitions between observed and gap states are modelled with distinct parameters. We denote the latter scenario as the deletion rate model (“+DEL”). Thus, the “+DEL” models have one additional free parameter when compared with global rate models.

### Simulation study

We used extensive simulations to explore the performance of the models introduced above. The length of the phenotypic sequences we simulated were purposely shorter than current molecular studies (Shendure et al. 2017) because empirical data sets are likely to contain relatively few “sites” in practice. For each combination of clade richness (50 or 100 species) and sequence length (10, 100, or 1000 positions) we produced 100 datasets under three modes of evolution: (1) a single rate shared by all sequence positions; (2) rates vary independent of position; and (3) rates are correlated between neighbouring positions. To simulate variation on the realized state space, both between and within sequences, we randomly sampled the dimension of the generating transition matrix (**Q**) for each sequence position from a common distribution. We fit the different models of sequence evolution to the replicates and compared support for each model across replicates using the Akaike Information Criteria (AIC).

We performed additional simulations to test the robustness of our analyses by replicating the observed characteristics of the *Gryllus* song sequences. We generated sequences under the homogeneous, gamma, and correlated models with 12 species, 728 sequence positions, and two possible states (i.e., sound or silence). We then aligned the sequences using the distance-based approach following the same procedure used for the empirical analyses (see *Supplementary material*). We added the MSA step here for the sole purpose of emulating the presence of gaps in the sequence as observed on the sequences used for the empirical data (see Figures 2 and 3). We fit the models of sequence evolution and compared model fit using AIC. See the *Supplementary material* for more information on all steps of the simulation study and the FigShare repository (*doi.org/10.6084/m9.figshare.c.4405310*) for the code and data to replicate analyses.

**Figure 1:**
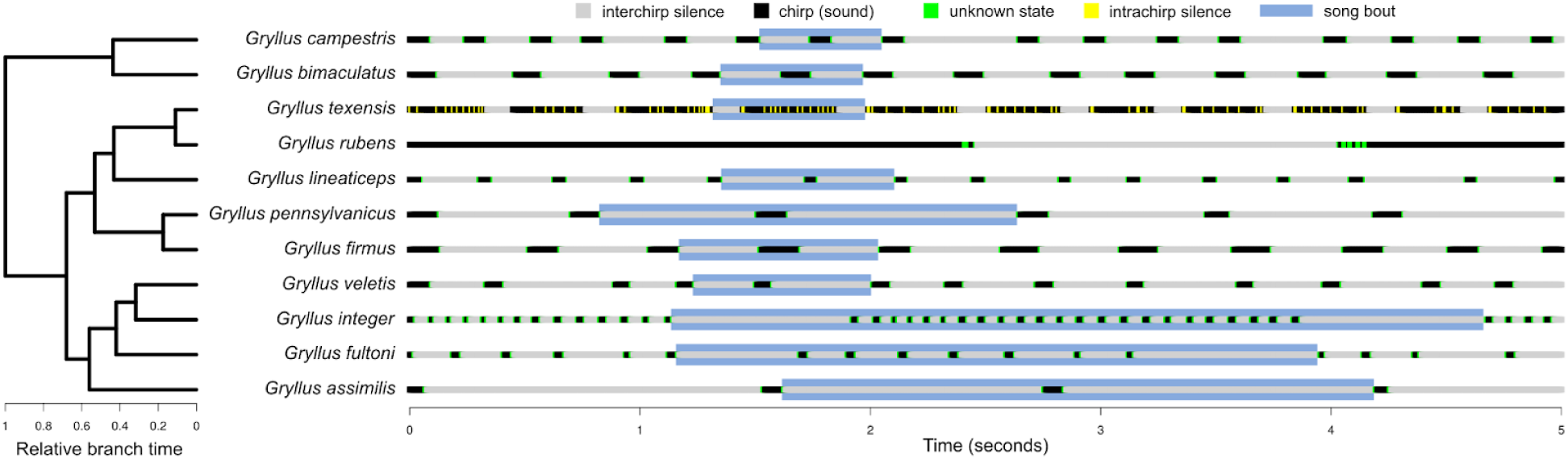
*Gryllus* male-display songs recorded over a 5s period as reported by Robillard et al. (2006). Sequences were rescaled from the original data to reflect the timing of the syllables (see *Supplementary material*). The maximum clade credibility tree on the left shows species relationships and relative branching times (see Figure S5). Length of song bouts vary among species and the light blue squares highlight the repeated patterns over time.

**Figure 2:**
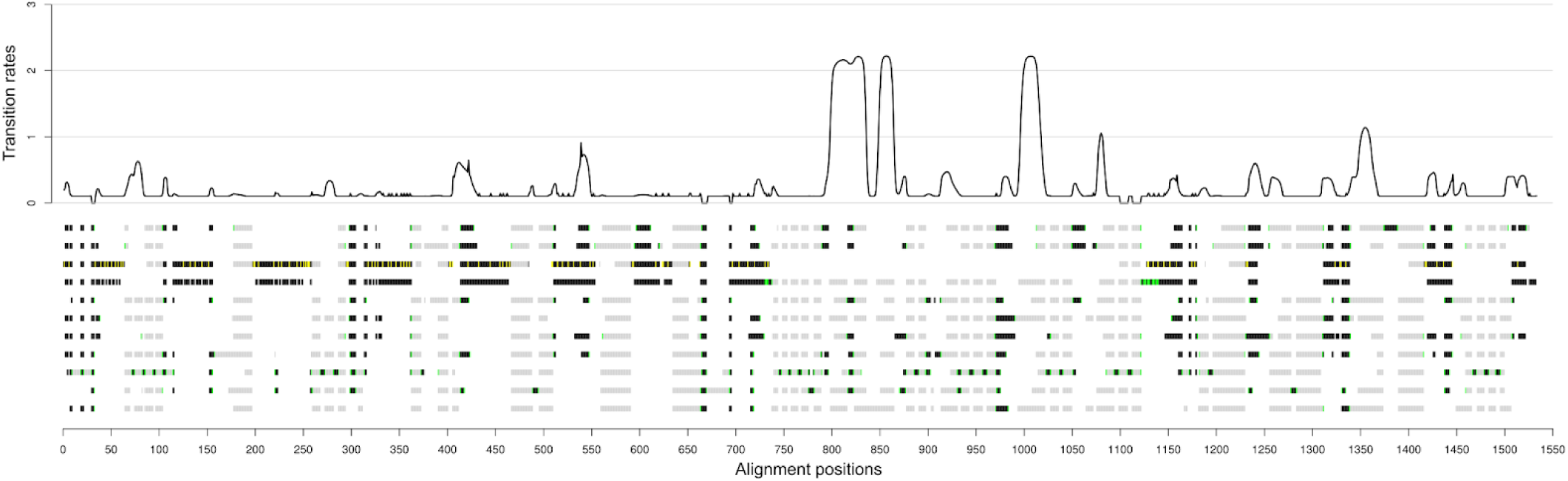
Model averaged transition rates estimated along the positions of the complete song sequence for 11 species of *Gryllus* (see Figure 1). Sequences show the distance-based alignment result based on the complete time-rescaled sequences and conditioned on the MCC tree. Order of species and color scheme is the same as shown in Figure 1. Gaps are represented as white spaces. Black solid line shows rates of transition between observed states (any color) and gaps (white). Red dots show rates of transition between observed states (estimated as zero for most positions).

**Figure 3:**
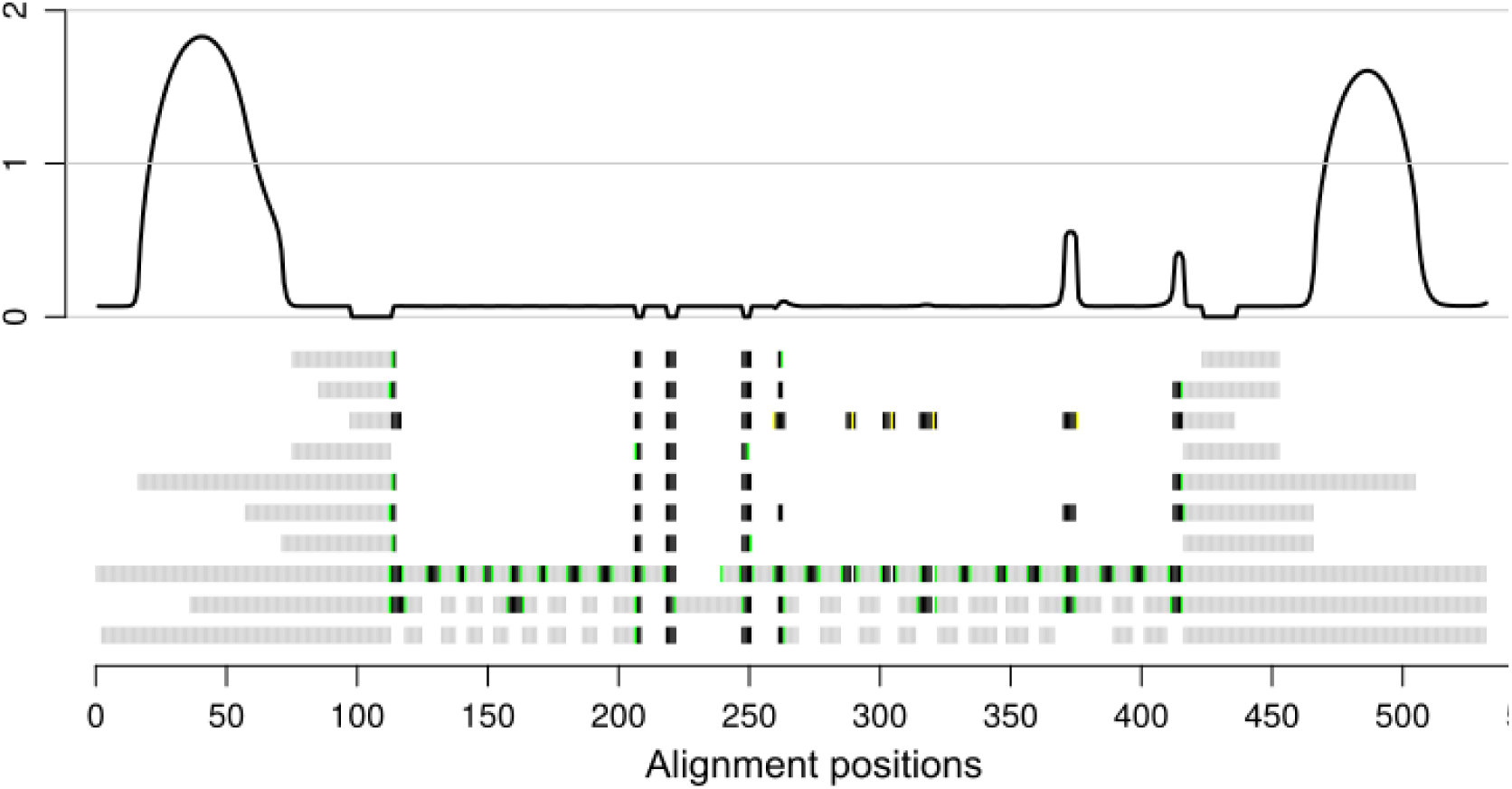
Model averaged transition rates estimated along the positions of song bouts extracted from the complete song recording of 10 species. Sequences show the distance-based alignment result based on a single bout sample extracted from the time-rescaled sequences and conditioned on the MCC tree (see *Supplementary material*). Order of species and color scheme is the same as shown in Figure 1 (except *Gryllus rubens* was not included). Gaps are represented as white spaces. Black solid line shows rates of transition for each sequence position.

### Analyzing cricket calling songs

We analyzed the evolution of male calling songs of nine North American and two European species of crickets (Grylloidea, *Gryllus*) to address two main questions: (1) Are rates of transition in the silent regions of the song (i.e., inter-chirp silence) distinct from rates estimated for chirp regions? and (2) Is there evidence for correlated rates of evolution across sequence positions? Male crickets produce sound by rubbing their raised forewings together and a sound syllable, or pulse, is produced during wing closure (Robillard et al. 2006), which alternates with periods of silence due to the time required to re-open the wings (i.e., intrachirp silence) as well as periods of inactivity (i.e., interchirp silence). Figure 1 shows the pattern of variation in the timing and length of interchirp silences and chirps for each studied species during an interval of five seconds derived from the data made available by Robillard and colleagues (2006). In their original study, Robillard et al. (2006) coded the movement of the wings (i.e., forewing opening and closure) as one character in a data matrix and then represented each interchirp silence by a number of characters proportional to its time duration (their “DO2” coding scheme). Here we conducted analyses based on their original data set (i.e., “DO2” coding) and also transformed the temporal pattern for each species into units of time such that each sequence position share the same time duration and every sequence has the same total length (Robillard pers. communication).

Since the sequences were recorded during a fixed time window of five seconds, the number of chirps present on a sequence varied among species (Figure 1). Here we deviated slightly from Robillard et al. (2006) and defined a song bout as a repeated pattern comprised by one chirp, or a group of chirps, and the adjacent pair of interchirp silences. Defined this way, a song bout is an adequate representation of a higher level organization of the phenotype and likely represents a more refined homology statement when compared to the whole sequence data. *Gryllus campestris*, for example, shows a short song bout (Figure 1 - highlighted in blue) with a single chirp separated by similar sized interchirp silences. On the other hand, *G. integer* shows a much longer song bout comprised by a group of chirps united by short length interchirp silences and separated by a pair of much longer interchirp silences (Figure 1 - highlighted in blue). We repeated these analyses by subsetting the data into song bouts. For this we extracted all potential bouts from the song sequence shown in Figure 1, producing a pool of song bouts for each species. We then constructed 100 data sets by randomly sampling a unique combination of song bouts for each species. The resulting collection of data sets represents variation in the song bout patterns for each species. Note that *G. integer* and *G. fultoni* are both represented by a single song bout (which was replicated in all data sets) and *G. rubens* was excluded from this analyses because, given our definition of song bout, the bout of this species was not completed within the five seconds period of recording.

We reconstructed the phylogeny of the *Gryllus* crickets in BEAST (version 2.5 - Bouckaert et al. 2014) using the same set of species and molecular data reported in Robillard et al. (2006) and citations therein (see *Supplementary Material* for more details). We conducted the alignment of the calling song sequences using both distance-based and model-based approaches, always conditioned on the maximum clade credibility tree (MCC) or a random sample from the posterior distribution from the BEAST analyses. We explored phylogenetic uncertainty by repeating the MSA conditioned on 100 randomly sampled trees from the posterior distribution using MAFFT (version 7.397, Katoh and Standley 2013) and a custom cost matrix (see *Supplementary Materials*). We then incorporated alignment uncertainty by randomly sampling 100 alignments proportional to their posterior probability using a model-based alignment approach as implemented in BAli-Phy and conditioned on the topology of the MCC tree (see *Supplementary Materials*). We repeated all alignments for each of the three alternative data sets — the original sequences (Robillard et al. 2006), the time-rescaled sequences, and the song bout data. We then fit models of homogeneous rates, independent rates among sequence positions (gamma model), and correlated rates among adjacent positions (correlated model). Finally, we compared model fit using the Akaike Information Criteria and performed model-averaging of rate estimates for each sequence position in function of the relative likelihood of each model (i.e., using AIC weights following Caetano et al. 2018).

## Results

### Performance of the models under simulation

Overall, our simulations showed an increase in the power to detect the generating model with increasing species number and sequence length (Figure S1). Comparing the distribution of ΔAIC per model across simulation replicates also suggested an increase in the potential to discern between models as sequence length increases and a lesser, but still detectable, effect due to clade richness (Figure S2). For very short sequences (10 positions), the homogeneous model performed well independent of the generating model, whereas for longer sequences (1000 positions), the generating model was almost always within the supported models (ΔAIC < 2). This result strongly suggests that more complex models are unlikely to be incorrectly favored due to problems with small sample size or elevated Type I error. Also, note that the correlated model was often supported when the generating model was correlated or homogeneous (Figure S2). We will discuss this pattern further below. Mean estimated parameter values were largely centered on the true parameter values independent of the size of the dataset or generating model, and the variance across replicates showed visible shrinkage in function of sequence length (Figure S3). Regardless, most cases resulted in less than 10% over or underestimation of rates (Figure S3).

It is important to note that the models fit herein are nested. For instance, the single rate model is a special case of the correlated model when the correlation equals 1 (i.e., a perfect correlation between each pair of sequence positions). We also noticed that high correlation values seemed to be associated with clusters of fast and/or slow rates along the sequence, as shown in Figure 3, and are likely associated with increased stability of rates within these clusters. The gamma rate model is also a special case of the correlated model when the correlation is 0, producing complete position-independent rates. We compared the distribution of rate estimates across sequence positions for each simulation replicate when the generating model was the single rate model (Figure S4) or the gamma model (Figure S5). The correlated model was able to recover a pattern of homogeneous rates along positions in most simulation replicates (Figure S4). In contrast, congruence between rates estimated by the correlated and gamma models are restricted to larger sample sizes (Figure S4). With a long sequence and a small phylogeny (1000 positions and 50 spp), rate estimates were similar between models, but they also showed very distinct AIC values. By contrast, when the correlated and gamma models were fit to our largest data set both the rate estimates and AIC values were similar between models.

**Figure 4:**
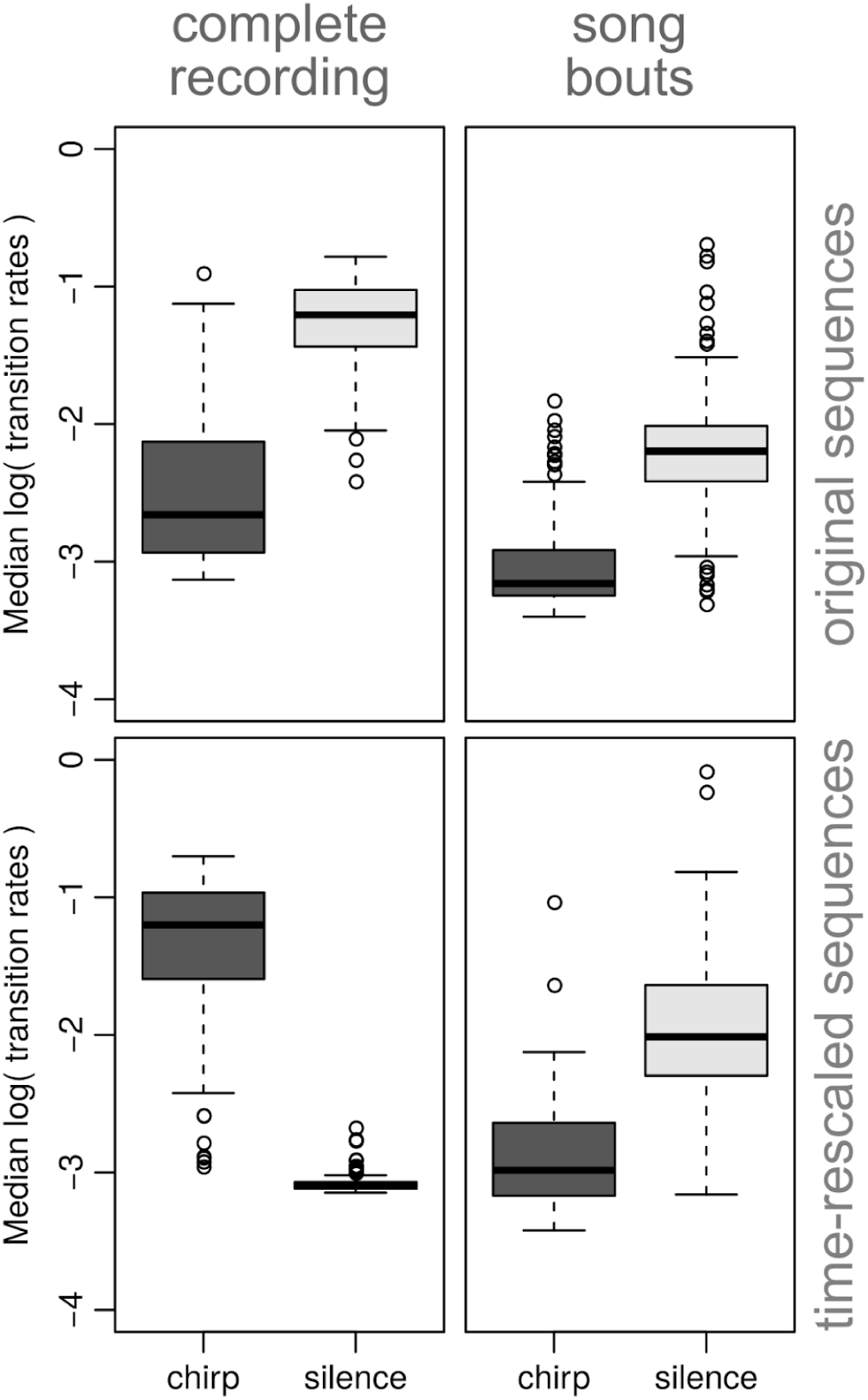
Distribution of median log-transformed rates of transition for chirp (sound) and silence (interchirp silence) regions. Top row results are based on original (DO2) sequences and bottom row results show time-rescaled sequences. Left and right columns show results with complete song sequences (5 seconds of recordings) and song bout sequences, respectively. Distributions of rates were computed across 100 randomly sampled alignments from the posterior distribution of the BAli-Phy analyses conditioned on the MCC tree and model-averaged in function of the relative likelihood of each model.

### *Gryllus* phylogeny and the alignment of phenotypic sequences

Our phylogenetic reconstruction based on molecular data produced a well-supported tree (Figures 1 and S7) with the same topology reported by Huang et al. (2000), which used a minimum evolution model (their Figure 3). Huang et al. (2000) found an incongruence in their maximum likelihood tree with *G. assimilis* appearing as sister to *G. fultoni*, which was not supported by our posterior distribution of trees (Figure S7). Our maximum clade credibility (MCC) tree supports a relationship with *G. assimilis* as sister to a clade comprised by *G. fultoni*, *G. integer*, and *G. veletis* (Figure S7).

We used the resulting MCC tree and a random sample of trees from the posterior distribution to perform alignments of the male calling song sequences (see examples in Figures 2 and 3). All BAli-Phy simulations converged within approximately 100,000 generations. The relatively short number of generations for convergence was likely due to the reduced dimensionality of the model, since we fixed many parameters by conditioning analyses on a fixed topology. We found a clear difference in the length of the distance-based and model-based alignments. MAFFT produced alignments consistently shorter than BAli-Phy, both when aligning the complete song sequences (Figures S8 and S10) and the song bout data (Figures S9 and S11). However, these approaches were conditioned on different phylogenetic trees and apply distinct models, thus a difference in alignment length is to be expected.

### Evolution of calling songs in crickets

We fit multiple models of sequence evolution to the male calling songs of *Gryllus* species while also exploring alternative data transformations, alignment approaches, and hypotheses of homology at distinct levels. Overall, the model of correlated rates of evolution was supported in all cases and was robust to multiple sources of uncertainty. Model averaged transition rates estimated for chirp and interchirp silence regions clearly showed faster rates of evolution are associated with silent regions of the calling songs in all but one case. We describe the results in more detail below.

When we performed distance-based alignments with the original song sequences (DO2; Robillard et al. 2006) conditioned on a random sample of 100 trees we found two supported models across replicates (Table 1). Both models incorporate correlated evolution between neighboring positions and differ only by a single parameter used to estimate separated transition rates between observed states and gaps (“+DEL” - Table 1 and Figure S12). The same analyses repeated with the time-rescaled sequences (see Figure 1) also found strong support for correlated rates (see Table 1 and Figure S8). We also found strong support for the correlated model for both the original and time-rescaled sequences after switching from distance-based alignments to a model-based approach (Table 1 and Figure S12). Finally, results were robust to distinct levels of homology. We found support for the same models of correlated evolution (see Tables 1 and 2 and Figure S13) even after changing the scope of our homology hypothesis from the complete 5 seconds period to only a single song bout (see Figure 1 for examples of song bouts). These results were also robust to variation in the song bout pattern as well as uncertainty associated with phylogeny and sequence alignment.

**Table 1:** Support for models of sequence evolution fitted to the complete song sequences (Figures 1 and 2) incorporating both alignment and phylogenetic uncertainty. Homo: single rate model; Gamma: independent rates fitted across sequence positions; Corr: correlated rates between neighboring positions. DEL: transition rates differ between observed states and gaps. Support was computed as the proportion of times each model showed relative Akaike Information Criterion (ΔAIC) and ΔAIC corrected for sample size (ΔAICc) lower than 2 units across a pool of 100 replicates incorporating either phylogenetic uncertainty (first 4 columns) or alignment uncertainty (last 4 columns). Column ‘AICw’ shows the median AIC weight for each of the models computed across replicates.

| Model | Distance-based alignment<br>(phylogenetic uncertainty) |  |  |  |  |  | Model-based alignment<br>(alignment uncertainty) |  |  |  |  |  |
| --- | --- | --- | --- | --- | --- | --- | --- | --- | --- | --- | --- | --- |
|  | Original data |  |  | Rescaled data |  |  | Original data |  |  | Rescaled data |  |  |
|  | AIC | AICc | AICw | AIC | AICc | AICw | AIC | AICc | AICw | AIC | AICc | AICw |
| Homo | 0.00 | 0.00 | 0.00 | 0.00 | 0.00 | 0.00 | 0.00 | 0.00 | 0.00 | 0.00 | 0.00 | 0.00 |
| Homo<br>+DEL | 0.00 | 0.00 | 0.00 | 0.00 | 0.00 | 0.00 | 0.00 | 0.00 | 0.00 | 0.00 | 0.00 | 0.00 |
| Gamma | 0.00 | 0.00 | 0.00 | 0.00 | 0.00 | 0.00 | 0.00 | 0.00 | 0.00 | 0.00 | 0.00 | 0.00 |
| Gamma<br>+DEL | 0.00 | 0.00 | 0.00 | 0.00 | 0.00 | 0.00 | 0.00 | 0.00 | 0.00 | 0.00 | 0.00 | 0.00 |
| Corr | **0.95** | **0.96** | **0.88** | 0.03 | 0.03 | 0.00 | **0.94** | **0.95** | **0.98** | **0.98** | **0.98** | **1.00** |
| Corr<br>+DEL | **0.32** | **0.17** | 0.12 | **0.99** | **0.99** | **1.00** | **0.30** | **0.15** | 0.02 | 0.04 | 0.02 | 0.00 |

**Table 2:** Support for models of sequence evolution fitted to song bout sequences (Figures 1 and 3) incorporating both alignment and phylogenetic uncertainty. Homo: single rate model; Gamma: independent rates fitted across sequence positions; Corr: correlated rates between neighboring positions. DEL: transition rates differ between observed states and gaps. Support was computed as the proportion of times each model showed relative Akaike Information Criterion (ΔAIC) and ΔAIC corrected for sample size (ΔAICc) lower than 2 units across a pool of 100 replicates incorporating either phylogenetic uncertainty (first 2 columns) or alignment uncertainty (last 2 columns). Column ‘AICw’ shows the median AIC weight for each of the models computed across replicates.

| Model | Distance-based alignment<br>(song bout uncertainty) |  |  | Model-based alignment<br>(alignment uncertainty) |  |  |
| --- | --- | --- | --- | --- | --- | --- |
|  | AIC | AICc | AICw | AIC | AICc | AICw |
| Homo | 0.00 | 0.00 | 0.00 | 0.00 | 0.00 | 0.00 |
| Homo+DEL | 0.00 | 0.00 | 0.00 | 0.00 | 0.00 | 0.00 |
| Gamma | 0.00 | 0.00 | 0.00 | 0.00 | 0.00 | 0.00 |
| Gamma+DEL | 0.00 | 0.00 | 0.00 | 0.00 | 0.00 | 0.00 |
| Corr | **0.83** | **0.87** | **0.63** | **0.76** | **0.81** | **0.64** |
| Corr+DEL | **0.75** | **0.48** | **0.37** | **0.69** | **0.49** | **0.36** |

We also assessed the performance of the model by comparing results using the empirical data against results from data sets simulated with the same dimensions of the observed data. Results based on simulated data sets showed support for the homogeneous rate model independent of the generating model (Table S2 and Figure S15) whereas the empirical data showed support for the correlated model (Tables 1 and 2). Note that these results differ from the simulation study described previously because here we performed sequence alignment with the simulated data in order to introduce alignment noise. The alignment step performed on simulated data re-organizes the positions and is likely to cause model misspecification. Nevertheless, the proportion of times the correlated model was supported when it was not the generating model was acceptably low (between 6 and 7% - see Figure S15), and indicate results reported here are not due to issues with elevated Type I error or small sample size.

We investigated rate variation along sequence positions by evaluating model-averaged rates along the sequence and testing for rate differences between chirp and interchirp silence regions of the alignment (see *Supplementary material* for more information). Figure 2 shows example of alignment and rates of transition estimated for the time-rescaled song sequences. Rates vary along the sequence and multiple regions emerge as being characterized by a sudden increase in transition rates. Rate estimates based on a distribution of phylogenetic trees (Figure S16) and a distribution of alignments (Figure S17) show similar patterns for both the original and time-rescaled data. However, the rate variation due to alignment uncertainty (Figure S17) seems to have a stronger effect than phylogenetic variation (Figure S16), likely a reflection of the stability of the *Gryllus* phylogeny (Figure S7). On the other hand, the pattern of rate variation estimated for the song bout data is very distinct (Figure 3), with higher transition rates restricted to interchirp silence regions located on the flanks of the bout sequences (Figures S18 and S19).

Finally, we computed the log-transformed medians (Figure 4) for the distribution of model-averaged rates along the song sequence grouped by chirp and interchirp silence regions. Then we performed a paired Wilcoxon signed-rank test to verify whether rate estimates are different between groups taking into account the effects of data transformation, distinct homology levels, as well as model uncertainty (see *Supplementary material*). Results show strong evidence for distinct transition rates between groups under all treatments (p < 0.001). Faster rates of transition were associated with silent regions of the calling song in most cases. However, the pattern is inverted when we used the whole song rescaled to time units (Figure 4, lower left).

## Discussion

### Model performance and identifiability

Here we fit models originally developed for molecular evolution (Yang 1993, 1995) to non-molecular sequences, which differ with respect to an unknown and variable state space among sequence positions and, importantly, reduced sample sizes. As far as we know, our work is the first attempt to expand the application of these models to the study of phenotypic evolution conditioned on phylogenies estimated from independent data. Fortunately, due to Yang’s elegant solutions (1993, 1995), the models implemented here are very flexible. These models vary between only one and four free parameters, which allowed us to describe distinct evolutionary modes for phenotypic sequences even when sample size could be a limiting factor.

We note that the minimal difference in number of parameters between models results in a small penalty applied to model comparison approaches. This can potentially lead to issues with model identifiability, especially when simpler models are nested within the more general model. This may sound alarming at first, but it serves to reinforce the argument that interpreting patterns informed by parameter values and investigating model behavior can be more informative than simply ranking models or rejecting null hypotheses (see discussion in Beaulieu and O’Meara 2016, Caetano and Harmon 2018, and Caetano et al. 2018). One might argue that the similar ΔAIC values for the single rate and correlated models shown on Figure S2 reflect issues with the power of the test. However, a plot of parameter values clearly shows that similar AIC values are due to equally well supported models. In other words, two distinct models point to the same pattern of evolution (Figure S4), thus accepting or rejecting the single rate model adds very little information to the interpretation of the data. When coupling model ranking with a thoughtful evaluation of parameter estimates, the models we present here are ideal for studying rates of evolution across the positions of non-molecular sequences, even in cases of small sample sizes. For instance, here we propose the use of model-averaged parameter estimates (following Caetano et al. 2018, and also see Beaulieu and O’Meara 2016) which is an ideal framework to shift the focus of the analyses from model ranking to rate estimates that incorporate uncertainty in model fit to the data.

### Patterns of evolution in cricket calling songs

We investigated the evolution of temporal patterns in male calling songs of 11 species of *Gryllus* using phylogenetic comparative models. We estimated the mode of evolution along positions of the song sequence which, unlike Robillard et al. (2006), relies on the implementation of explicit statistical models. We found strong support for autocorrelation of rates of evolution along the sequences. This result corroborates previous hypotheses that the temporal pattern of calling songs evolve through the expansion and contraction of both chirp and interchirp silence regions by insertions and deletions of pulses and the elements of a chirp (Otte 1992, Robillard et al. 2006, Robillard and Desutter-Grandcolas 2011). Evidence for autocorrelation of rates also support the scenario of context-dependent rates of evolution where similar regions of the sequence are more likely to share a common rate.

Our comparison of model-averaged rates of evolution between chirp and interchirp silence regions resulted in mixed results when contrasting the whole sequence and the song bout data, but were otherwise robust to the variation derived from both phylogeny and alignment uncertainty (Figures 4 and S20). For instance, results from whole sequences showed higher rates for either chirp or interchirp silence regions, depending if either the original or the time-rescaled data was used (Figure 4 - top row). On the other hand, results from sequences restricted to a single song bout per species were congruent between the original and rescaled data, both showing evidence for faster rates of transition associated with interchirp silence regions (Figure 4, bottom row).

One potential issue with the results obtained from the whole sequence data is that repetitions are a significant challenge for sequence alignment approaches, because it is not clear to which of the copies a sequence segment is supposed to align (Notredame 2002; see also Desutter-Grandcolas and Robillard 2003, Robillard et al. 2006). For instance, the alignment in Figure 2 shows different histories estimated for the multiple song bouts represented within each of the sequences whereas Figure 3 shows a much more conservative alignment solution. Is it possible that higher rates of evolution estimated for chirp regions when using the whole song recording might be associated with the misalignment of duplicated patterns. Another important factor is that the original data, as coded by Robillard et al. (2006), define the sequence units as to-and-fro movements of the forewings, which constitute both the behavioral homology and the sound unit of the calling songs. Thus, the incongruent results between the whole sequences rescaled to time and the other treatments suggest that the behavioral units constitute a stronger homology statement than time patterns and better reflect the evolutionary units for this multidimensional trait (also see Robillard et al. 2006 on this point).

Knowledge about the genetic architecture of temporal patterns in cricket calling songs is very limited, but evidence shows the number of pulses per second, one of the components of the song, is associated with a polygenic architecture (Blankers et al. 2018 and citations therein). Thus, the large scale temporal patterns studied here very likely have a genetic basis at least as complex. Moreover, many sources of selection are known to act upon calling songs, since these serve as species recognition traits (Otte 1992, Izzo and Gray 2004, Gray et al. 2016), are under sexual selection (Forrest 1983, Hedrick 1986, Zuk et al. 2008), and can attract unwanted attention in the form of male competitors (Jang 2011) or predators (Hedrick 2000). More specifically, female preference generally selects for longer song bouts (Hedrick 1986), but response to selection is counterbalanced by the risk of predation (Hedrick 2000). Our findings that interchirp silence regions evolve faster are therefore interesting because multiple selective pressures acting on a multidimensional trait can limit the phenotypic space that lineages can explore. However, changes associated with silent regions might allow phenotypic diversification on a separated temporal pattern axis which might be under relatively lower evolutionary constraints.

### Current challenges and moving forward

Our main objective was to evaluate approaches originally designed for molecular data when applied to phenotypic sequences, since these share fundamental characteristics. We found that the most challenging step was the construction of a multisequence alignment. First, different from molecular data, there is no generalized homology statement applicable across any phenotypic sequence. Non-molecular sequences require careful definition of *a priori* homology hypotheses, especially when the trait under study is comprised of repeated patterns. This contrasts with molecular studies that utilize homology statements based on sequence similarity defined with generic pipelines (Pearson, 2013). The amount of subjectivity introduced in the analyzes will, of course, vary with the type of data. For instance, when studying development sequences each stage serves as both a relative position and special quality criteria for homology (Remane 1952, Mindell and Meyer 2001), whereas the data used herein was amenable to different levels of homology. Second, virtually all alignment software assume molecular sequence data as both the input format and for the alignment search scheme (e.g., BAli-Phy; Suchard and Redelings 2006). Few approaches, such as MAFFT, are flexible enough to allow the use of any sequence of interest under custom searching parameters. Finally, transition rate models need to make explicit assumptions about the meaning of gap states. Here we translated gap states into absence of the trait at the sequence position (i.e., “indel”), although, it is possible for alternative translations depending on the data type and question.

## Conclusion

Here we expanded models originally described for molecular data to ask questions about evolutionary rates of phenotypic sequences using phylogenetic trees. We implemented models to fit a single rate across the sequence, independent rates for each sequence position, and autocorrelated rates of evolution. We showed that these models perform surprisingly well even when sample sizes are orders of magnitude smaller than average molecular sequences. These results pave the way for the investigation of a large series of data types. In fact, these models can be applied to any multidimensional trait ordered in a sequential structure.

The diversity of characteristics used in comparative studies is remarkably low when compared to the breadth of phenotypic variation known to occur across the tree of life. Our view of macroevolution is restricted to a few types of phenotypes (mostly morphology) evaluated one at a time on phylogenetic trees of few well-known clades whereas, in reality, traits are much more diverse and co-evolve under varying degrees of organization. We hope our study encourages the community to explore a larger variety of traits, especially overlooked phenotypes such as behavioral displays and developmental series. Many important technological advances to data collection and management, such as miniaturization of recording devices, increased Internet bandwidth, and Terabyte-size storage, can help drive a revolution in the number and variety of traits explored in comparative studies. Unfortunately, phenotypic evolution studies of most clades are lagging behind due to an inordinate focus of research resources toward charismatic (“model”) groups and translational research. A change in pace sorely depends on increased support for natural history, taxonomy, and biodiversity studies, especially of understudied clades.

## Acknowledgments

We thank T. Navok, J. Boyko, and M. Araya-Salas for their insightful critiques and constructive comments on an earlier version of this manuscript. We especially thank T. Robillard for important information about the data and *Gryllus* biology. We thank B. Redelings for help with BAli-Phy and K. Katoh for help with MAFFT and insightful comments on sequence alignment. DSC would also like to thanks the Coordenação de Aperfeiçoamento de Pessoal de Nível Superior (CAPES: 1093/12-6) for partial support on this project.

## Supplementary material for

### 1) Simulation results - Figures and tables

**Figure S1:**
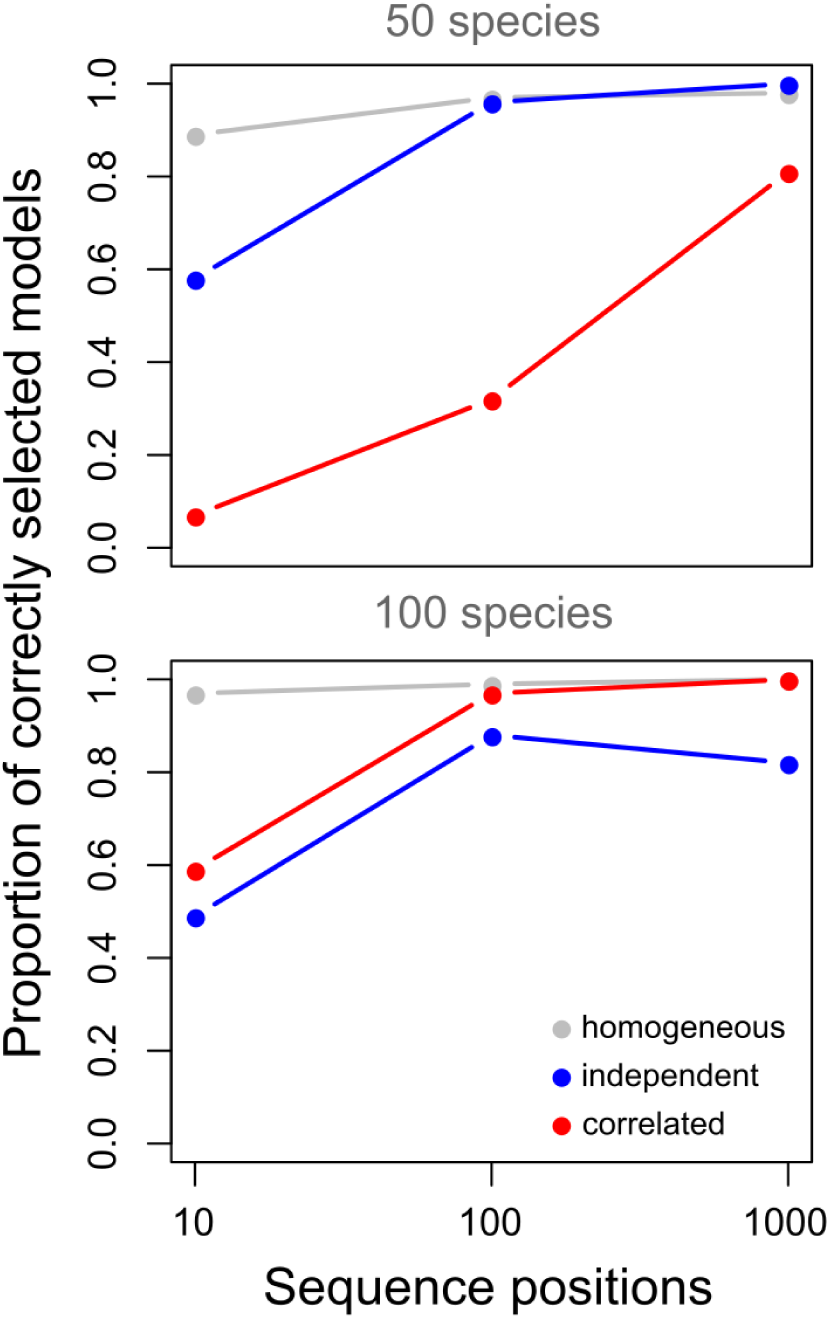
Results from 100 simulation replicates under each of the generating models. Gray lines show results for the homogeneous rate model. Blue lines show results for the independent rates (Gamma) model. Red lines show results for the correlated model. Top plot shows results with 50 species and bottom plot with 100 species. The proportion of correctly selected models was computed by counting the number of times, across the 100 replicates, that the generating model showed ΔAIC < 2.

**Figure S2:**
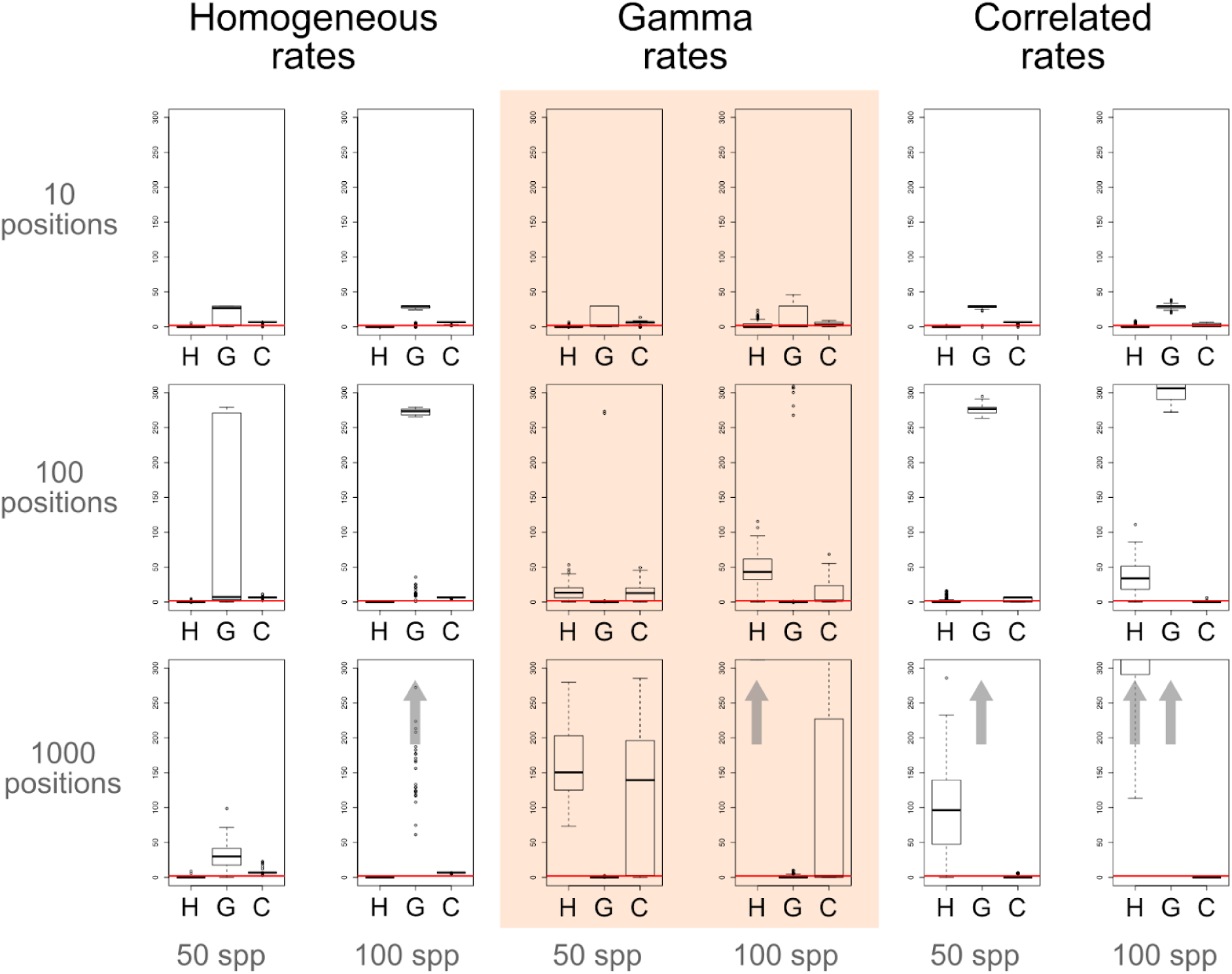
Distribution of pairwise ΔAIC for each of the models across simulation treatments. Each plot show results for a window from 0 to 300 ΔAIC units. Any result with mean value above 300 ΔAIC units was omitted from the plot and represented by a grey arrow. The salmon square in the middle highlights the results for the independent rates (Gamma) model. To the left are results for the homogeneous model and to the right the results for the correlated rates model. Top row shows results for simulations with only 10 positions, middle row with 100 positions and bottom row with 1000 positions. Each column separates the simulations between phylogenetic trees with 50 and 100 species. The red lines mark the ΔAIC < 2 threshold which groups the models favored by the data.

**Figure S3:**
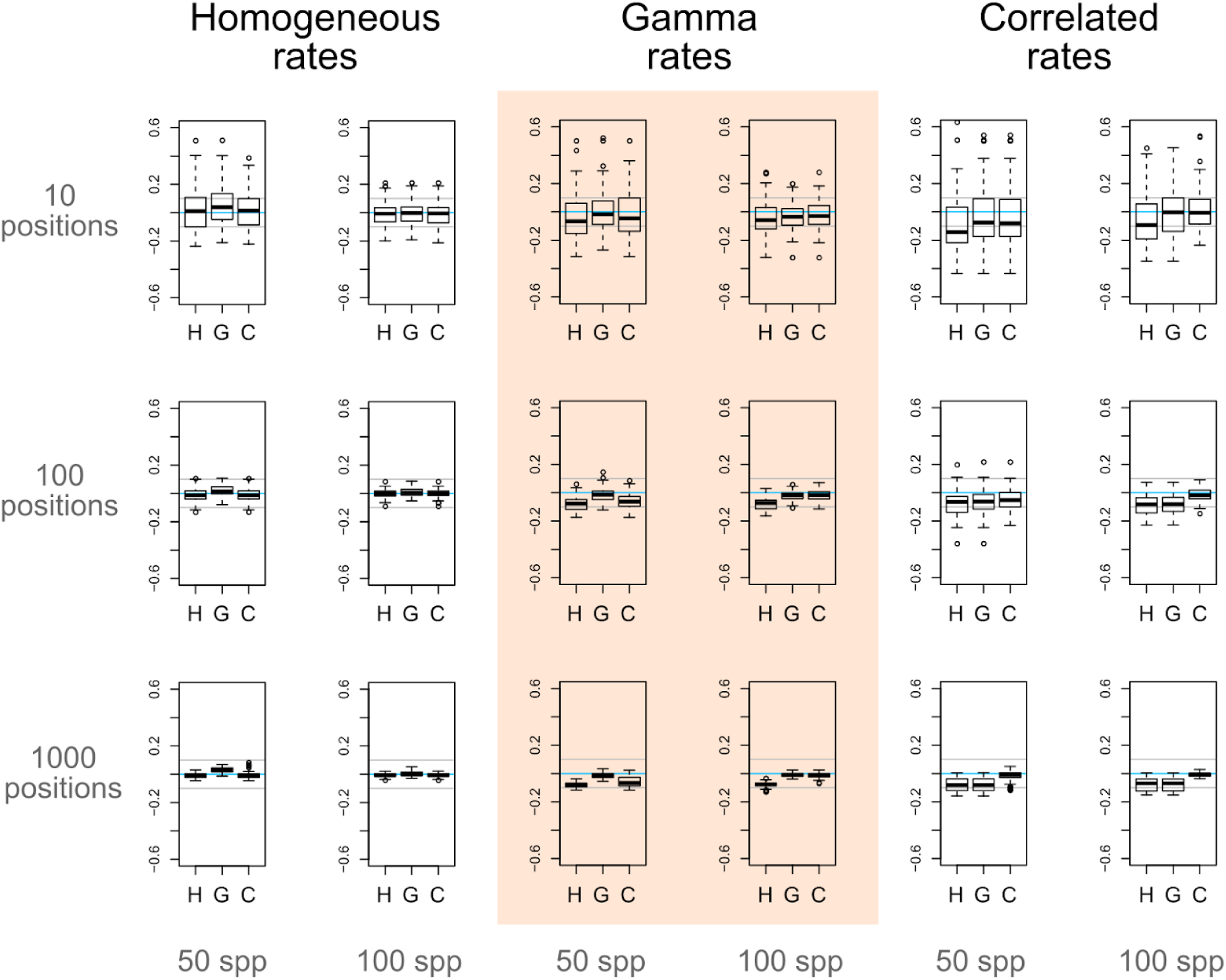
Distribution of parameter estimates for each of the models across simulation treatments showed as the relative average distance (computed as proportion of the true parameter) from the true rates of evolution for each sequence position. Light blue lines mark the 0 point (i.e., no difference in average rates). Grey lines mark the 10% deviance limit above or below the true rates. The salmon square highlights the results for the independent rates (Gamma) model. To the left are results for the homogeneous model and to the right the results for the correlated rates model. Top row shows results for simulations with only 10 positions, middle row with 100 positions and bottom row with 1000 positions. Each column separates the simulations between phylogenetic trees with 50 and 100 species.

**Figure S4:**
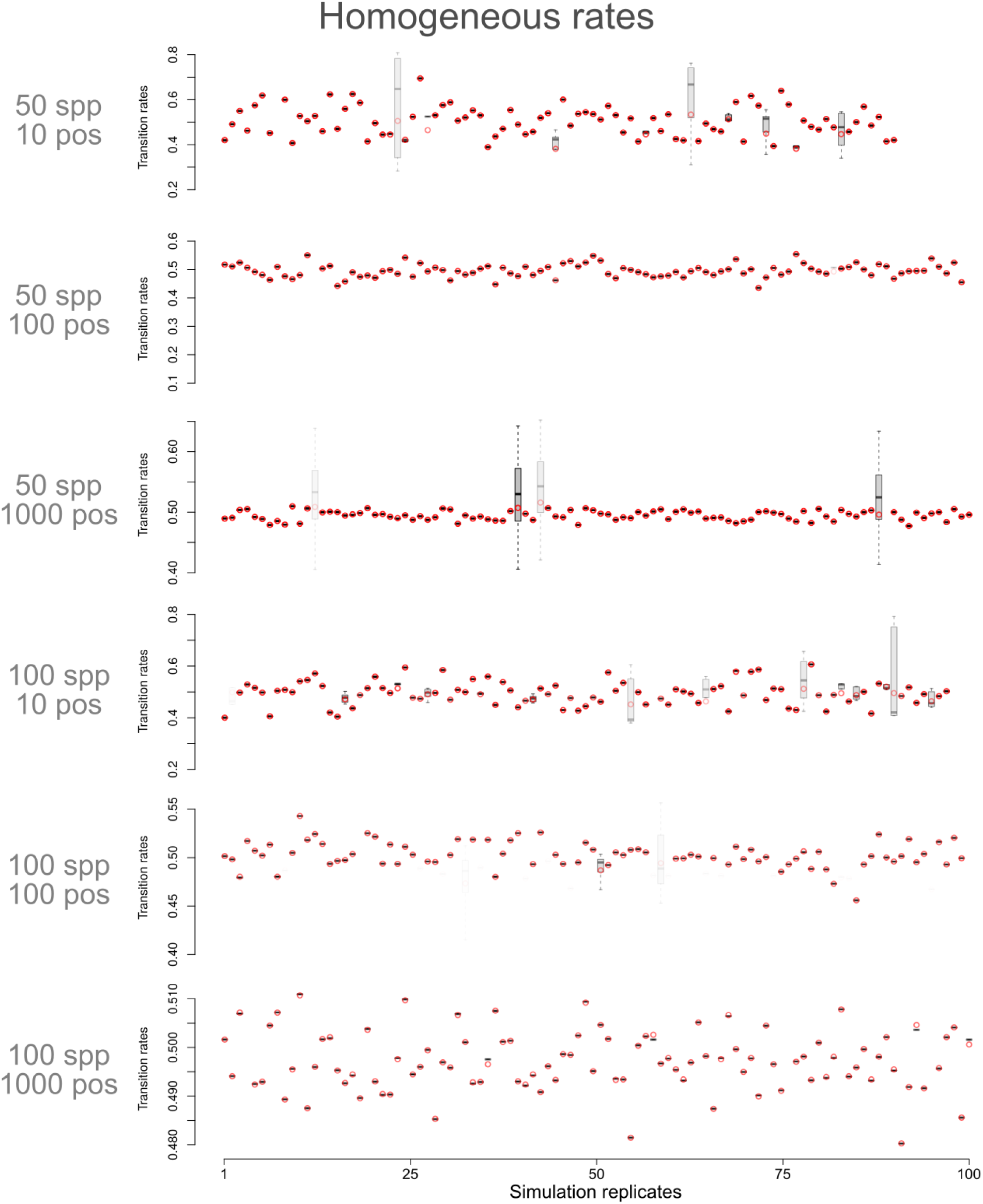
Distribution of rate estimates for the single rate model (red circles) and correlated rates model (overlaid box plots) for datasets simulated under a single rate model. Each plot shows results for a different simulation treatment varying the number of species (spp) and the number of sequence positions (pos). The x-axis of the plots show results for the 100 simulations replicates for each of the treatments (order is arbitrary). Red circles are transition rates under the single rate model for each simulation. Rates under the correlated model are shown as box plots and black horizontal lines represent box plots with no variation (i.e., cases in which the correlated model recovered a single rate across the sequence). The transparency of the color for each simulation point (red circle + box plot pair) is proportional to the ΔAIC between models. In other words, opaque symbols show models with similar AIC and transparent symbols represent models with distinct AIC scores.

**Figure S5:**
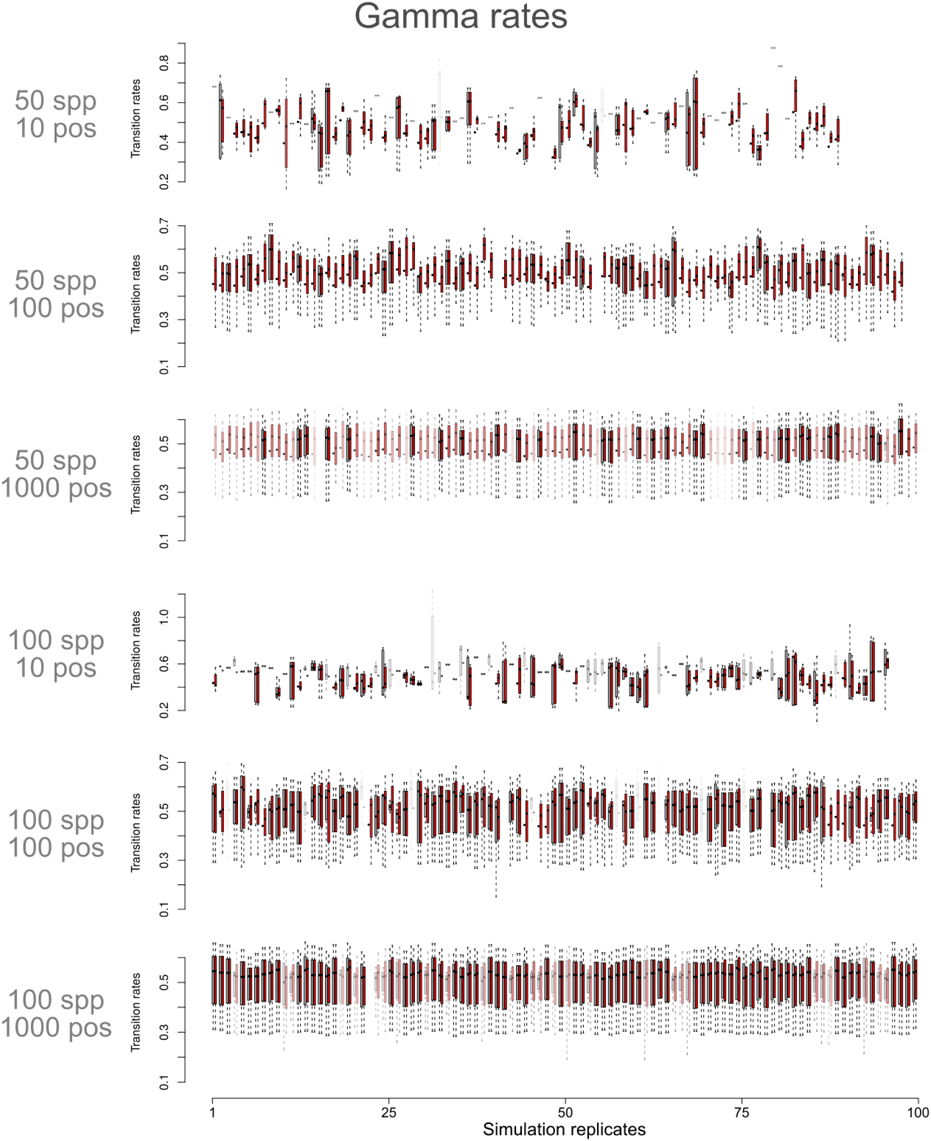
Distribution of rate estimates for the gamma model (red box plots) and correlated model (grey box plots) for datasets simulated with rate heterogeneity along sequence positions (i.e., with Gamma distributed rate categories). Each plot shows results for a different simulation treatment varying the number of species (spp) and the number of sequence positions (pos). The x-axis of the plots show results for the 100 simulations replicates for each of the treatments (order is arbitrary). Each pair of red and grey box plots represents the distribution of rates under the gamma and correlated model, respectively. Each pair of box plots (denoting results from the same simulation replicate) is separated by a small white space. The transparency of the color for each simulation (red + grey box plot pair) is proportional to the ΔAIC between models. In other words, opaque symbols show models with similar AIC and transparent symbols represent models with distinct AIC scores.

## 2) Empirical case: Evolution of calling song in Gryllus crickets

### 2.1) Estimating the Gryllus phylogeny from molecular data

We downloaded sequences for mitochondrial 16S and cytochrome B (cytb) from GenBank following accession numbers reported by Robillard et al. (2006 - see their Table 1). We aligned the 16S and cytb sequences using MAFFT (Katoh et al. 2013) and fitted molecular evolution models for each gene using jModelTest 2 (Darriba et al. 2012). Results showed support for the GTR+I+G model for both genes under AIC and BIC model selection criteria.

In order to infer a tree with ultrametric branch lengths and incorporate uncertainty in tree estimation we used BEAST2 (Bouckaert et al. 2014) with a birth and death tree prior and relative branching times. Our initial analyses using default priors showed that transition rates between C and G as well as G and T were near 0. This creates issues with the mixing of the chain because the Gamma distribution does not allow rates to be 0. In other words, the Markov-chain Monte Carlo (MCMC) simulation starts to sample too close to the hard bound of the prior distribution. For instance, the initial MCMC returned an Effective Sample Size (ESS) of less than 10 units for these transition rates.

We updated our selection of priors on the transition rates to a uniform distribution with range informed by the results from our preliminary analysis with the same data (see Figure S7). Initial values for the MCMC chain were set to the mean of the posterior distribution for each of the parameters estimated using the initial default priors. The range of the improper uniform prior distributions were set to be proportional to the range of the posterior distribution under BEAST2 default priors. Then, we ran the MCMC for 10 Million generations and excluded 25% of the samples as burn-in. We checked the trace of the posterior, likelihood, and priors to check for convergence of the chain as well as the ESS of each parameter of the model to check for good mixing. We also computed the ESS for the tree topology using the R package “rwty” (Warren 2017).

The resulting phylogenetic tree showed good posterior support, with the majority of nodes showing posterior probabilities above 0.9 (Figure S8). When compared with the trees reported by Robillard et al. (2006), the topology of the Maximum Clade Credibility (MCC) tree is closer to the phylogeny using molecular data and acoustic characters (their figure Figure 2A). The maximum clade credibility tree reported on Figure S8 is congruent with results from Huang et al. (2000).

**Figure S6:**
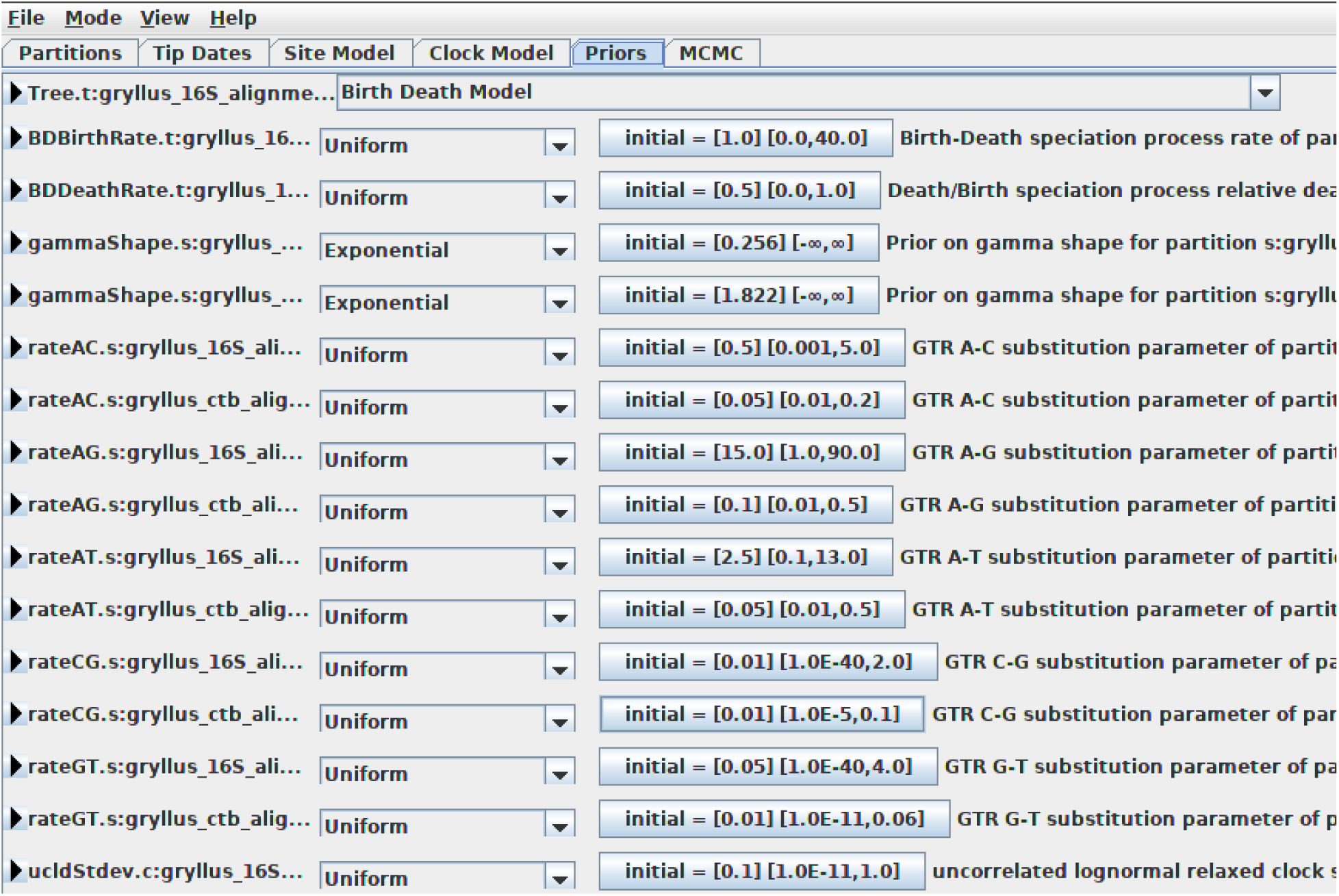
Starting values and prior distributions for the parameters of the BEAST2 analyses using 16S and cytb sequences for *Gryllus* crickets. Figure is a screen capture from the BEAUti user interface set to the “Priors” tab.

**Figure S7:**
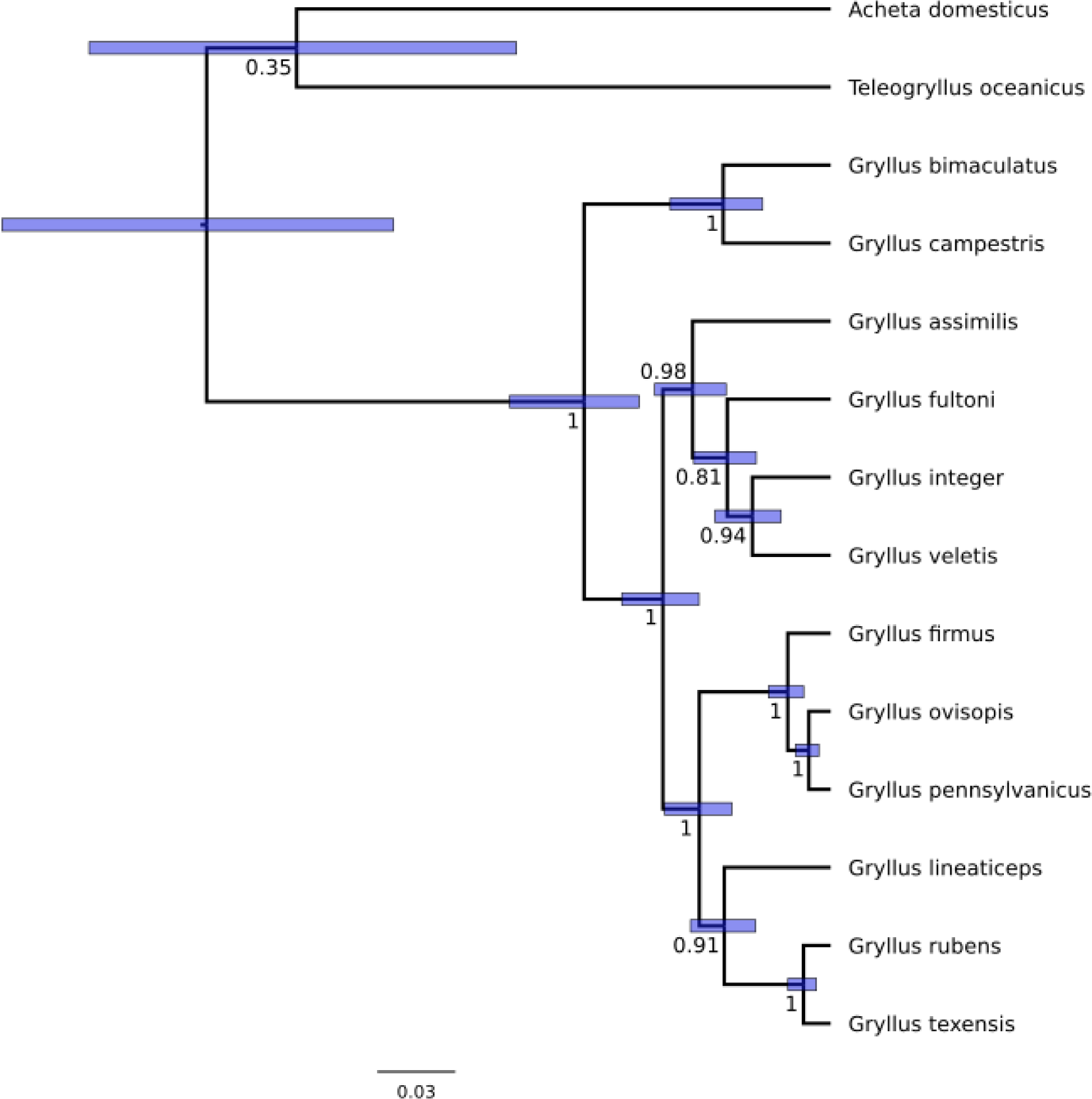
Maximum Clade Credibility (MCC) tree. Numbers at the nodes show posterior node support computed as the proportion of times each node was observed across the posterior samples. Purple bars show the 95% credible interval for the node heights in units of relative branching times.

### 2.2) Rescaling cricket songs sequences to units of time

Robillard and colleagues (2006) coded the temporal pattern of the male *Gryllus* calling songs using a single character state (coded as A) for each syllable, or pulse. Then they implemented two approaches to code the silent regions. First they used a single character (coded as G) to denote a silent region independent of its length (i.e., DO1 coding). Alternatively, the authors used a string of silent characters (i.e., a string of G) to code the silent segment with a number of characters equal to the time interval divided by the syllable period (i.e., DO2 coding). Here we used only the DO2 character coding in our analyses because, as discussed by Robillard et al. (2006), the DO2 code scheme takes into account the time information of the interchirp regions and performs better than the DO1 coding scheme. We refer to the DO2 coding as “original sequences” in the main text.

Each song sequence was recorded for an interval of 5 seconds. Under the DO2 coding, the sequence for a species has a number of characters equal to 5 seconds divided by the time period of a single pulse. In other words, the sequences reported by Robillard et al. (2006) are in units of pulse. Since the time period of a pulse varies between species, the unaligned sequence lengths also vary. Here we explored the temporal pattern of the male calling songs by transforming the sequences from units of pulse back to units of time (seconds). For this we computed the time period for the pulse of each species and rescaled the sequences such that each sequence would have the same length. Thus, in the time rescaled data reported here, a sound character is not equivalent to a pulse, but it is a representation of a common unit of time between the species. We report results with both type of sequence for each analyses conducted in this study.

### 2.3) Selecting song bout sequences

The calling song of male *Gryllus* crickets is a complex trait emerging from repeated pulses separated by periods of silence. A pulse is a complete movement of the wings, which is the homologous behavior that produces the sound (Otte 1992, Robillard et al. 2006). When pulses are repeated in a tight sequence, with no intervals (or intervals of very short length) a chirp (or trill) is produced (see Otte 1992). The silent interval between chirps (or trills) is called interchirp silence. Here we define a song bout as a higher level repeated pattern for which a group of chirps (or trills) and interchirp silences constitute the units of repetition. Figure 1 shows the whole 5 seconds of song recordings for each of the species of *Gryllus* included in this study. The majority of species shows a simple song bout, comprised by a single chirp (or trill) separated by an interchirp silence region which repeats multiple times through the recorded sequence. Species *G. campestris*, *G. bimaculatus*, *G. texensis*, *G. lineaticeps*, *G. firmus*, *G. veletis*, and *G. assimilis* belong to this group. In contrast, *G. integer* and *G. fultoni* have more complex song bouts. Figure 1 shows that, for both species, there is a region of short chirps connected by short interchirp silences which is flanked by much longer interchirp silences. We defined these regions as the song bouts. Finally, it is not possible to evaluate the song bout pattern of *G. rubens* because it’s longer song bout was not fully recorded within the 5 seconds window. In other words, there is no evidence of a higher level repeated pattern on the song sequence recorded for this species. As a result, the song sequence of *G. rubens* was excluded from all analyses based on song bouts.

We manually separated each song bout from the complete song sequences for all species. This generated a file with multiple song bouts per species, except for *G. integer* and *G. fultoni* whose sequences only presented a single bout. Then, in order to include within sequence variance in song bouts, we produced data sets by randomly drawing one song bout per species. We were able to produce 100 <u>unique data sets</u> by mixing song bouts of all species.

### 2.4) Alignment of cricket songs using MAFFT

The default algorithm in MAFFT (Katoh et al. 2013) produces an alignment based on a guide tree, which is inferred from the sequences as part of the algorithm using different approaches. MAFFT is a distance-based alignment approach that, by default, applies the standard BLOSUM62 score matrix (Henikoff and Henikoff 1992). In contrast, there is no standard cost matrix for the alignment of non-molecular sequences.

The *Gryllus* male calling song sequences used in the present study were coded with four states (following Robillard et al. 2006): sound characters (A), interchirp silence (G), and intrachirp silence (C). Robillard and colleagues (2006) also used the states R and M to indicate polymorphic states: either sound or intrachirp silence (R) and either sound or interchirp silence (M). In order to align these sequences we used the option ‘`--textmatrix`’ implemented in MAFFT (supported on versions ≥ 7.120 - Katoh et al. 2013). This option allows the user to define a custom scoring matrix, which substitutes the default score matrix. In this matrix, scores > 1.0 indicate an important pairing and scores between 0.0 and 1.0 are set for different elements that share similarities (with larger values carrying more weight than lower values). Finally, mismatches between important and dissimilar elements are set to negative values. With this scoring scheme it is possible to use the alignment algorithm in MAFFT without the need to assume patterns associated with molecular evolution processes.

Table S2 shows the custom scores used in this study. We set cost penalties such that matches between sound elements would be the most informative, followed by matches between sound elements and the R states (either sound or intrachirp silence). Then, we set matches between either intra- or interchirp silences as slightly less informative than sound elements. Further combinations were set following these same general rules.

We repeated the alignment procedure for the complete song sequences with 100 phylogenetic trees randomly sampled from the posterior distribution of trees inferred using MCMC in order to incorporate the effect of uncertainty in topology and branch lengths. In the case of the song bouts, we repeated the alignment across 100 data sets incorporating variance on the song bout expression across the recorded sequences and conditioned on the MCC tree. The alignment with the original sequences showed a distribution of sequence length varying between 1151 and 1245 positions (Figure S9A). In contrast, the alignment using the time rescaled sequences based on the same sample of trees had a minimum of 1501 and maximum of 1717 positions (Figure S9B). Finally, the alignment using the song bout sequences varied between 335 and 365 positions for the original sequences and 500 and 600 positions for the time-rescaled sequences (Figure S10).

**Table S1:** Custom score matrix used to perform the alignment of *Gryllus* song sequences in MAFFT. Column ‘Pair code’ shows the HEX code required by MAFFT to identify character symbols. Column ‘Pair states’ shows the correspondent pair of states. Finally, column ‘Score’ shows the relative score for matches and mismatches.

| Pair code | Pair states | Score |
| --- | --- | --- |
| 0x41 0x41 | A A | 1.5 |
| 0x41 0x52 | A R | 1.3 |
| 0x52 0x41 | R A | 1.3 |
| 0x47 0x47 | G G | 1 |
| 0x43 0x43 | C C | 1 |
| 0x47 0x52 | G R | 0.7 |
| 0x52 0x47 | R G | 0.7 |
| 0x43 0x52 | C R | 0.6 |
| 0x52 0x43 | R C | 0.6 |
| 0x43 0x47 | C G | 0 |
| 0x47 0x43 | G C | 0 |
| 0x41 0x47 | A G | - 0.5 |
| 0x47 0x41 | G A | - 0.5 |
| 0x41 0x43 | A C | - 0.5 |
| 0x43 0x41 | C A | - 0.5 |

**Figure S8:**
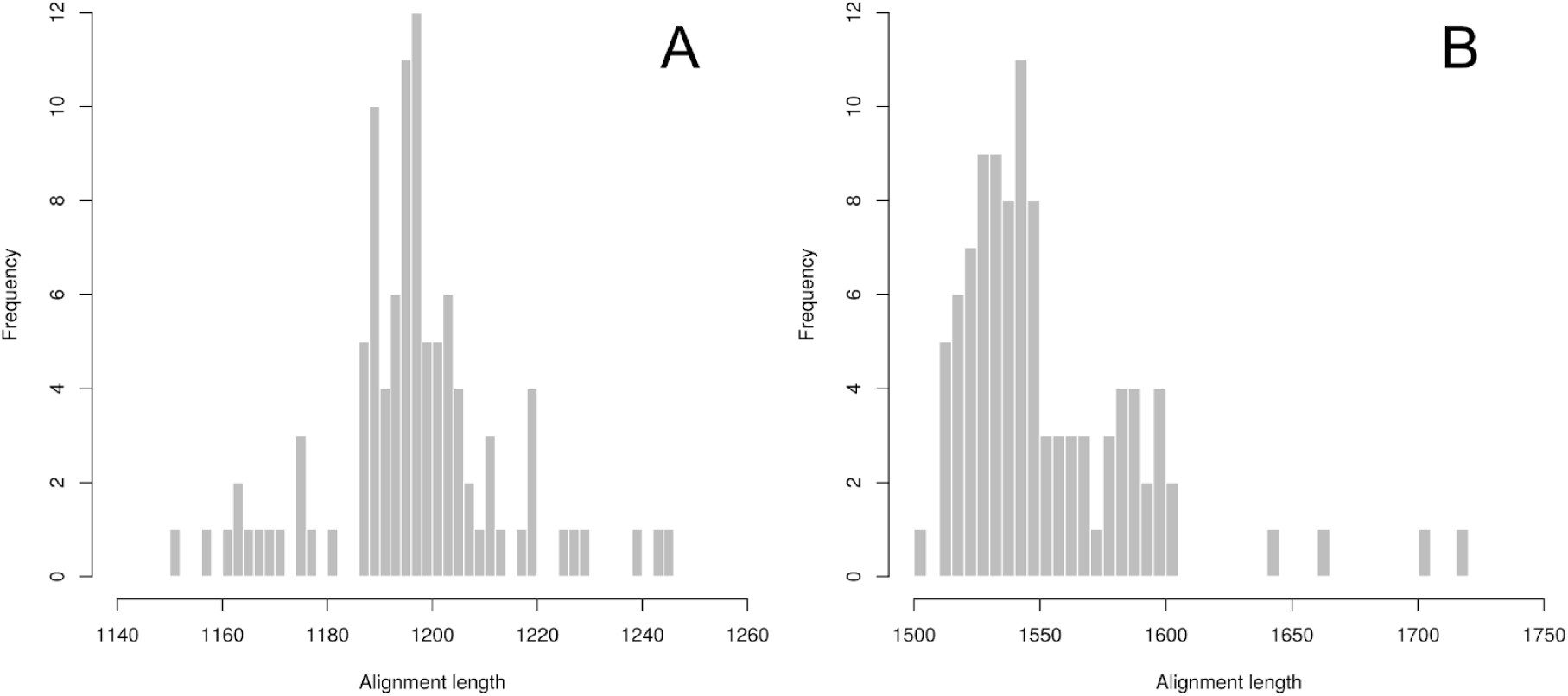
Distribution of alignment lengths for complete song sequences using MAFFT over a sample of 100 trees from the posterior distribution. A) Original sequences (DO2). B) Time rescaled sequences.

**Figure S9:**
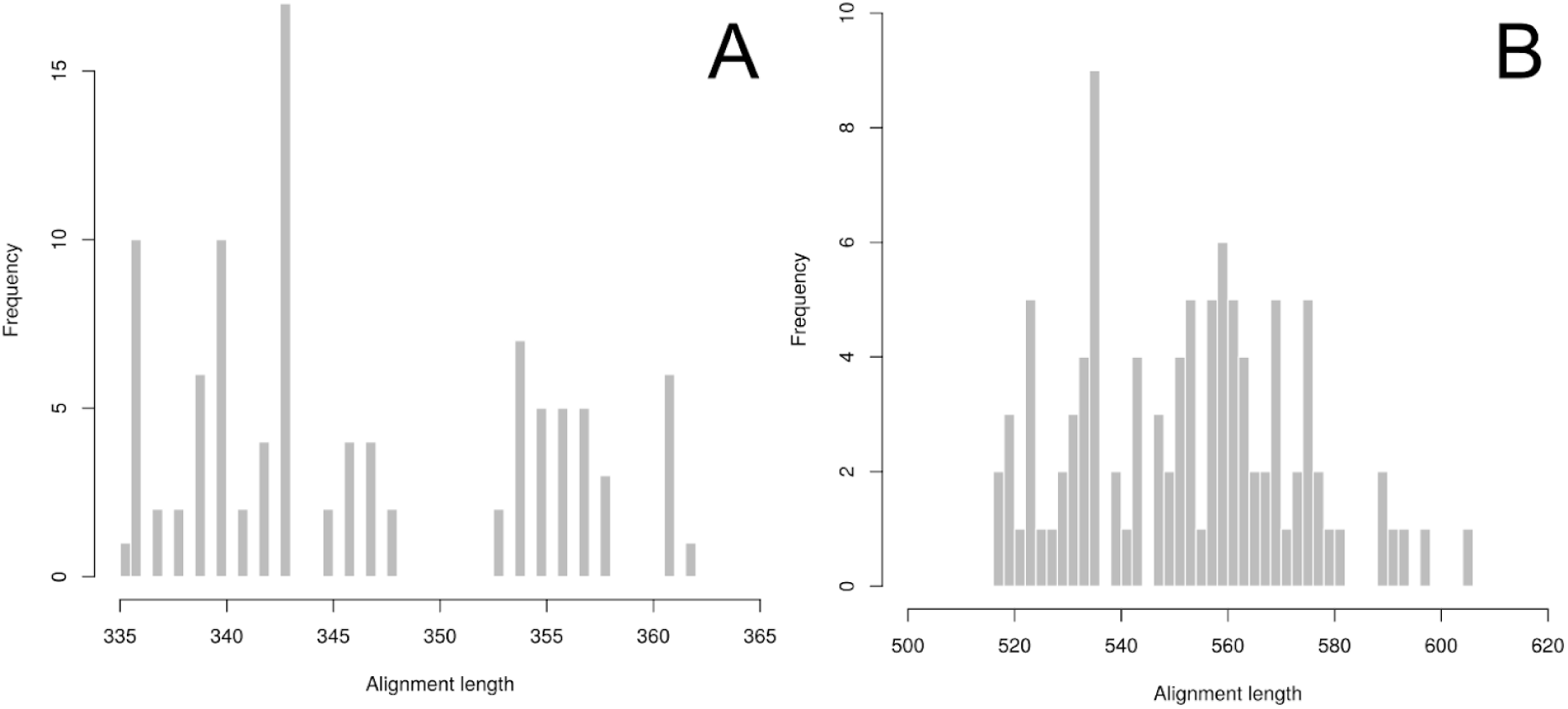
Distribution of alignment lengths for the song bout data. A sample of 100 data sets was produced by random sub-sample of song bouts from the complete song sequences. Distance-based alignments were conducted using MAFFT conditioned on the MCC tree. A) Bouts from original sequences (DO2). B) Bouts from time-rescaled sequences.

### 2.5) Statistical alignment of cricket songs using BAli-Phy

We also performed alignment estimates using BAli-Phy in order to incorporate alignment uncertainty based on a statistical alignment approach conditioned on the Maximum Clade Credibility (MCC) tree. For this we constrained the topology to be the same as the MCC tree, but allowed branch lengths to vary because BAli-Phy estimates branch lengths following the parameters of the model of sequence evolution (Redelings and Suchard 2005). For this we set the following options:

~~~
# Disable moves on the topology
disable = topology
# Set the phylogeny
tree = Gryllus_only_tree_MCC.tre
~~~

We ran two independent MCMC chains for each data set (DO2 and time rescaled) and evaluated convergence by checking trace plots of prior and posterior probabilities on Tracer (Rambaut 2018). We also checked if the effective sample size for the parameters were reasonably high (> 500). We computed the Gelman and Rubin’s potential scale reduction factor (Gelman and Rubin 1992) for all parameters as implemented in the function `gelman.diag` of the R package “coda” (version 0.19.2, Plummer 2006) for each parameter of the model.

The alignment with the original sequences (Robillard et al. 2006), using BAli-Phy and conditioned on the topology of the MCC tree (Figure S11) showed length varying between 1409 and 1727 positions (Figure S11A). In contrast, the alignment using the time rescaled sequences resulted in a posterior distribution of alignments much longer than the one based on the original sequences (minimum of 2145 and maximum of 2445 positions) (Figure S11B). Finally, the distribution of alignments using the song bout data conditioned on the topology of the MCC tree resulted in alignments between 435 and 626 positions for the original sequences and 650 and 900 positions for the time-rescaled sequences (Figure S12).

**Figure S10:**
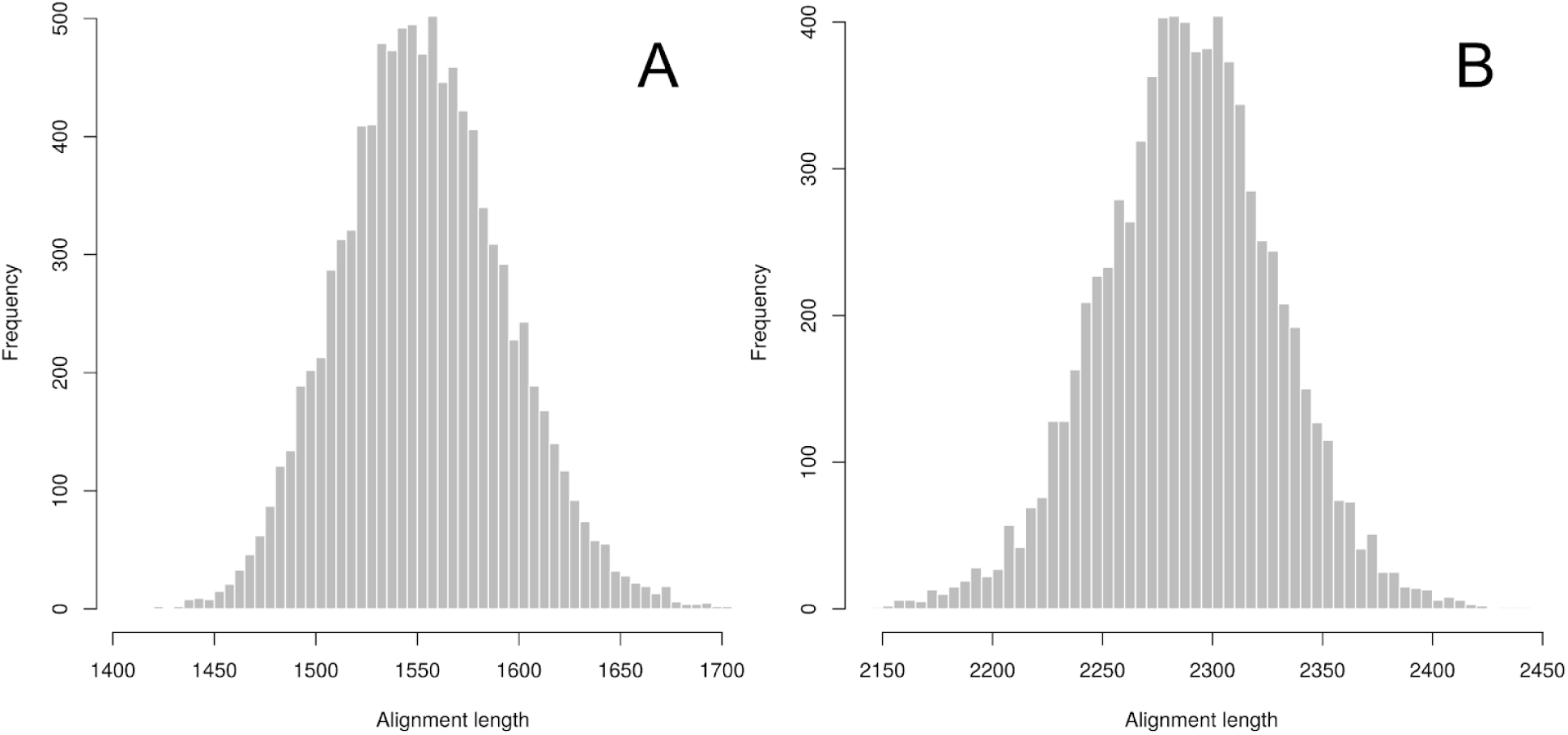
Distribution of alignment lengths for the complete song sequences from the posterior distribution of BAli-Phy conditioned on the topology of the MCC tree. A) Original sequences (DO2). B) Time rescaled sequences.

**Figure S11:**
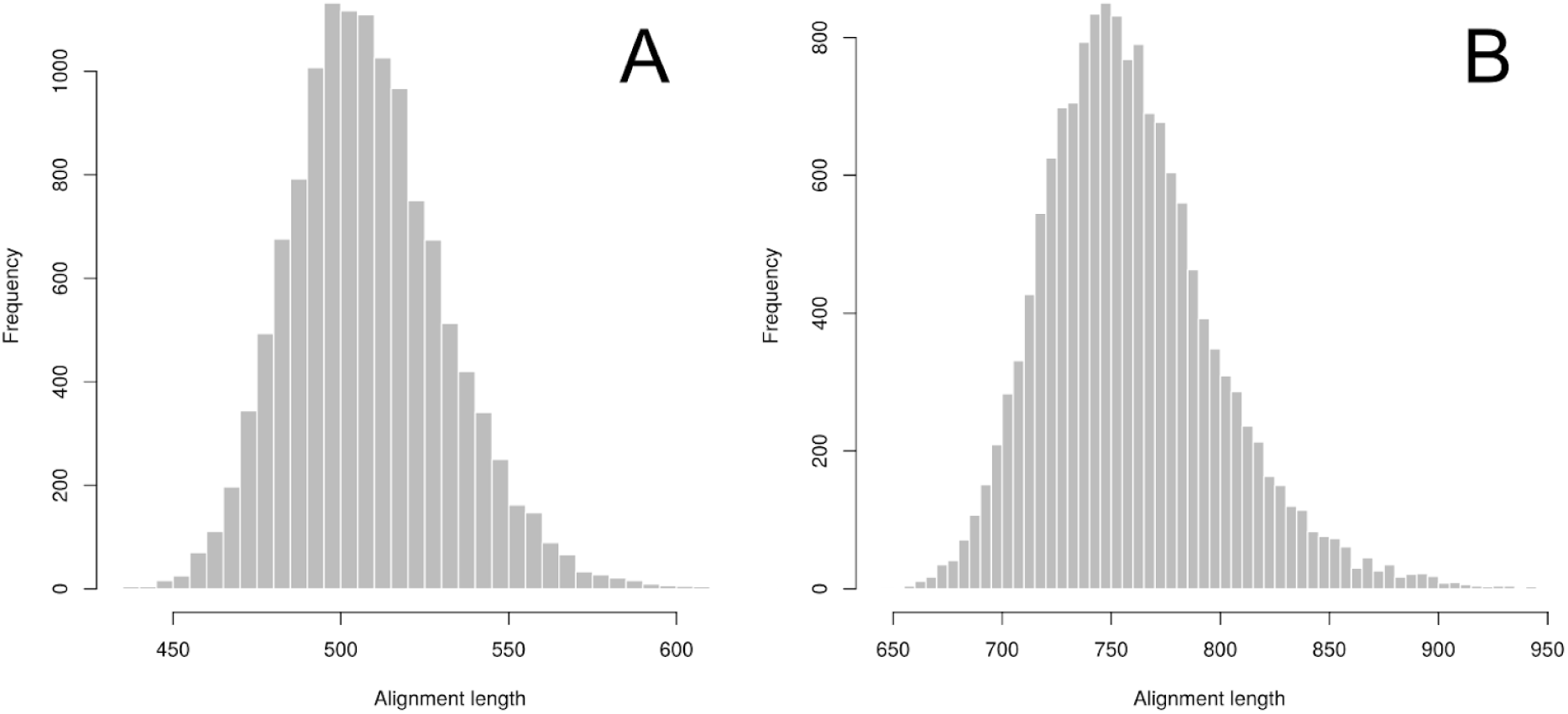
Posterior distribution of alignment lengths for the song bout data from BAli-Phy. A) Bouts from original sequences (DO2). B) Bouts from time-rescaled sequences.

### 2.6) Support for alternative models

The plot below lists the delta AIC values for the fitted models across all data transformations and alignment approaches implemented on this study. These complement results reported in Table 1 (main text).

**Figure S12:**
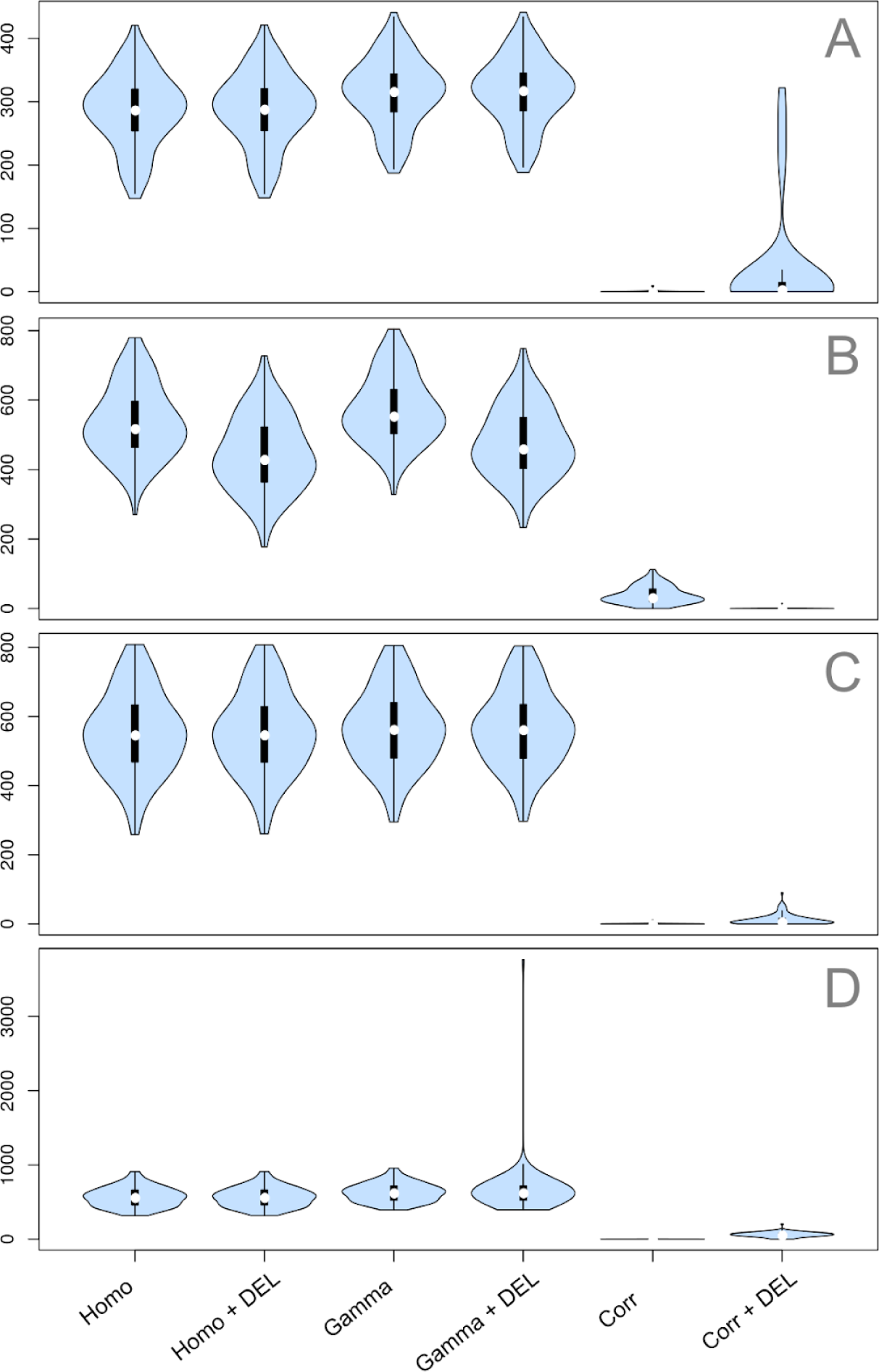
Distribution of delta Akaike Information Criterion (ΔAIC) for models of sequence evolution applied to both original (DO2) and time rescaled song sequences. A) Original sequences and distance-based alignments. B) Rescaled sequences and distance-based alignments. C) Original sequences and model-based alignments. D) Rescaled sequences and model-based alignments. Distributions on A and B were estimated across a random sample of 100 trees. Distributions on C and D were estimated across a random sample of 100 alignments from the posterior distribution of alignments conditioned on the topology of the maximum clade credibility tree (MCC). See Table 1 for a description of the models.

**Figure S13:**
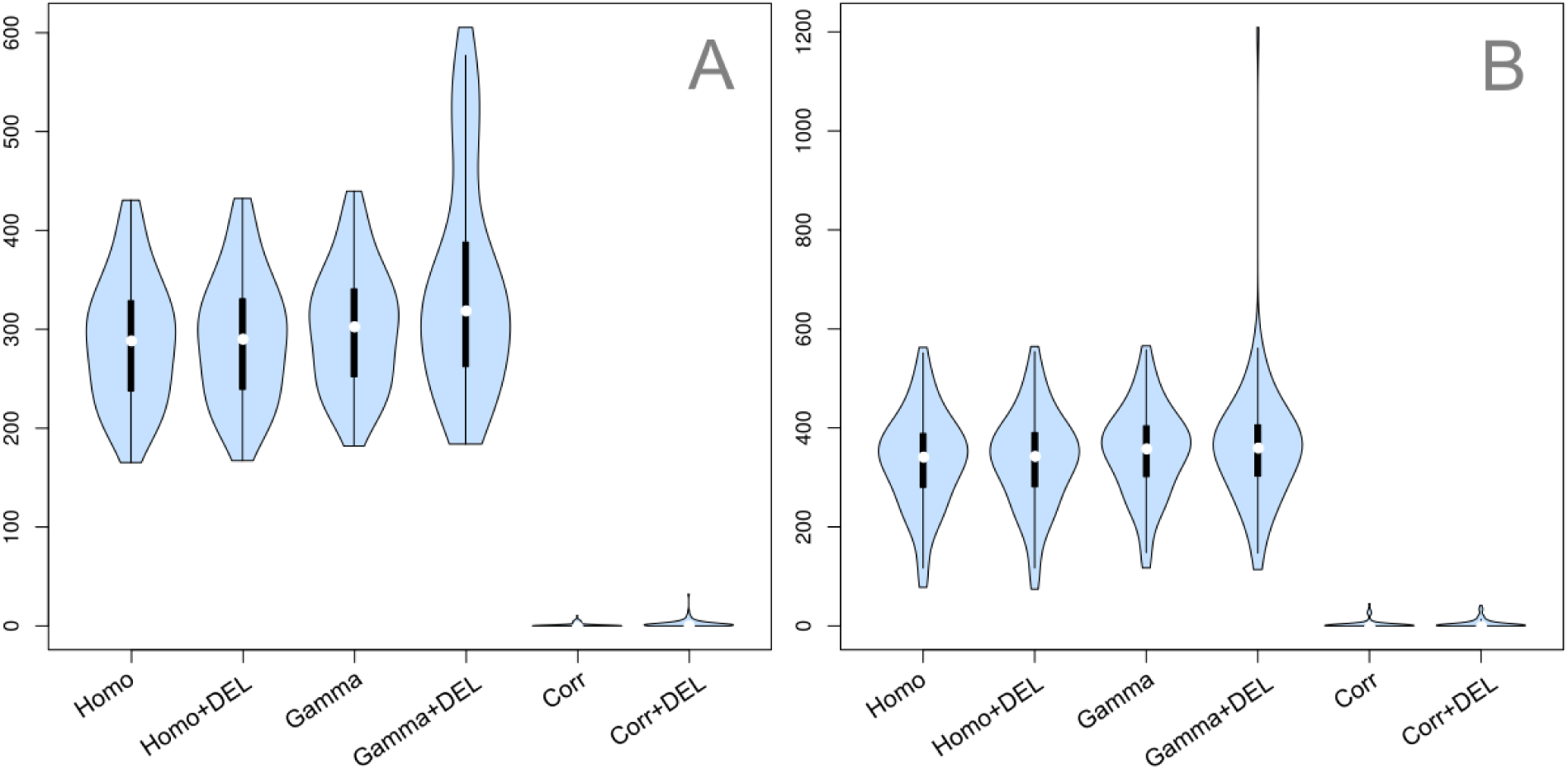
Distribution of delta Akaike Information Criterion (ΔAIC) for each model of sequence evolution applied to a single song bout for each species of *Gryllus* (see Figure 3 of the main text). A) Results based on distance-based alignments conditioned on a sample of 100 trees. B) Results with a sample of alignments from the posterior distribution of BAli-Phy conditioned on the Maximum Clade Credibility tree (MCC). See Table 2 and main text for a description of the models.

### 2.7) Testing the robustness of results

In order to test the robustness of our results we simulated datasets with the same dimension of the *Gryllus* male calling song sequence data. We generated 100 ultrametric phylogenies with 12 extant species and tree height of 1 time unit. We used independent Markov models with the same parameters values we chose for the extended simulation study to generate phenotypic sequences with 728 positions for each phylogeny---same as the alignment of the observed data. We did not replicate the simulations using the original sequence data (Robillard et al. 2006) because it is not straightforward to simulate sequence traits of different length between species. We set the number of states to be fixed and equal to 2 across sequence positions---the state “A” represents sound elements and “G” silence elements.

We performed sequence alignment with the distance-based approach as implemented in MAFFT and using the same score matrix shown in Table S1. We did not align the sequence using BAli-Phy because of the significant computing resources necessary for the analyses coupled with the lack of a built-in signal to stop the MCMC sampler at a pre-selected number of MCMC generations. These characteristics of BAli-Phy generate non-trivial problems to the automatization of the pipeline across a large sample of data sets. For instance, one could use a parallel running program in order to automatically verify a stopping condition given the data on BAli-Phy output files and halt the BAli-Phy run by pairing the correct running process. However, such program is beyond the objectives of the present study. Finally, we conducted model fit and model comparison using the Akaike Information Criterion.

The distribution of alignment lengths for the simulated data is similar to the empirical results (Figure S15), meaning that we can safely compare results of the simulation with the empirical data. The proportion of times that the correlated model was selected when it was not the generating model is acceptable (Figure S16 and Table S3). In contrast, the power of the test is relatively low, with only about 20% of the simulation replicates returning the correct model when the correlated model was the generating model. The low Type I error recovered by these simulations means that the chance the *Gryllus* sequences returned spurious evidence for correlated rates of evolution among sequence positions is acceptably low. Thus, we are confident about the results despite the low power due to the reduced size of the data set. Please refer to our discussion on the main text.

**Figure S14:**
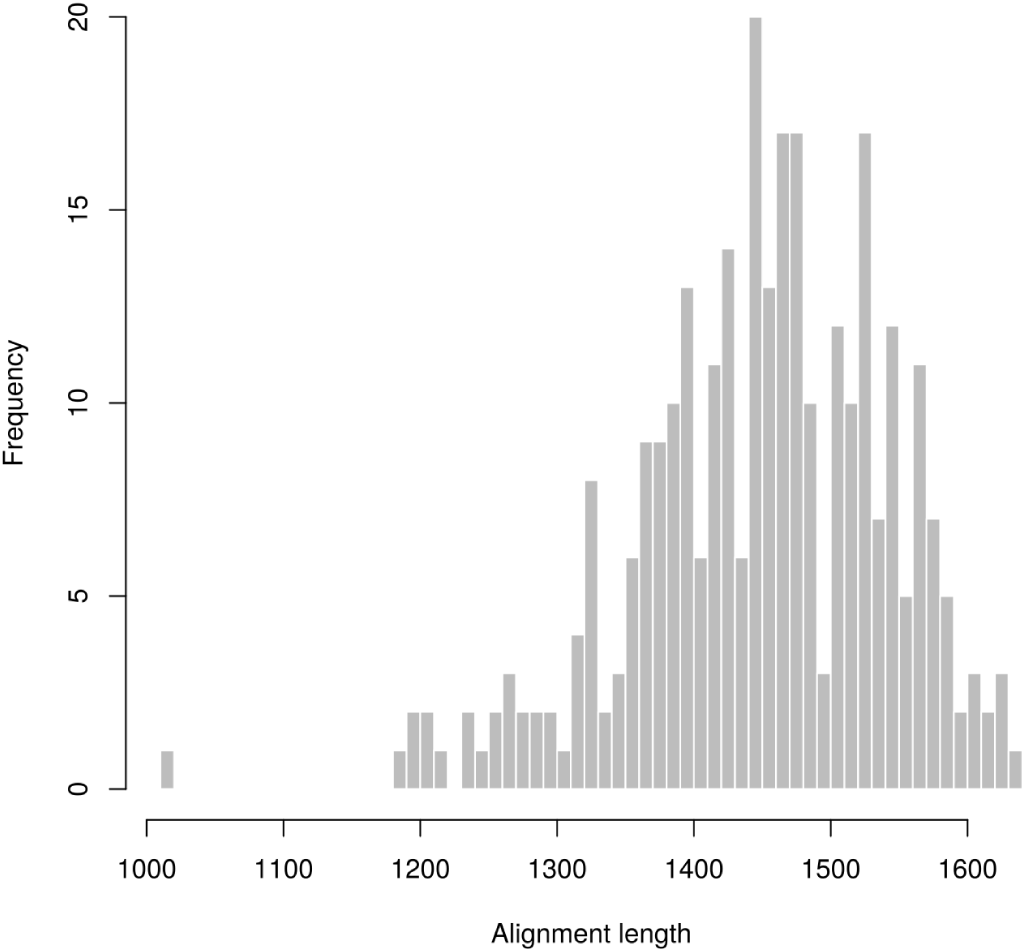
Distribution of alignment length for sequences simulated with the same dimension of the empirical complete song sequence data (12 species and 728 positions).

**Figure S15:**
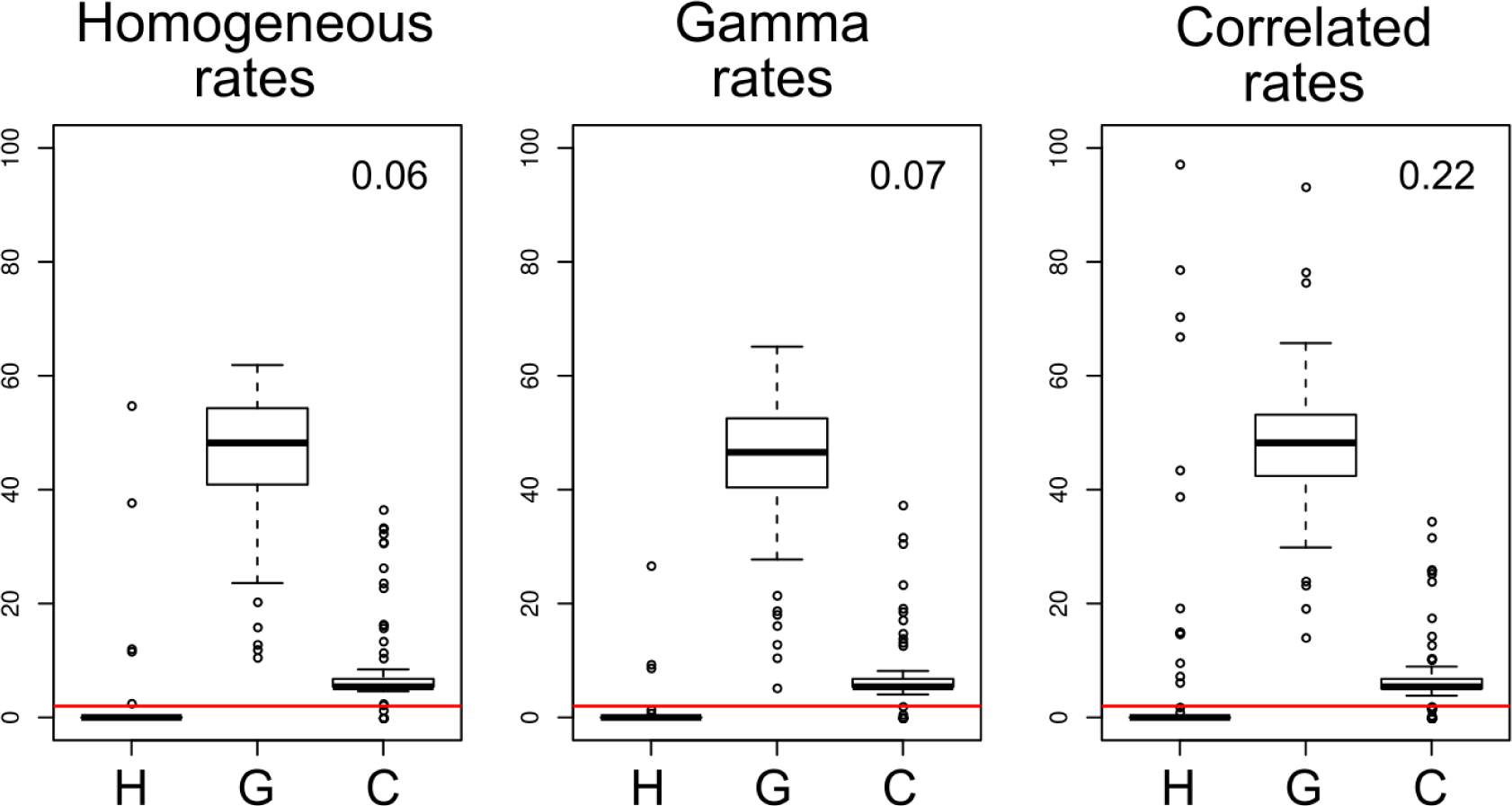
Distribution of pairwise ΔAIC across 100 simulation replicates for each of the three generating models with data sets showing the same number of species and sequence positions of the empirical data. H: homogeneous rates model; G: independent (Gamma) rates model; C: correlated rates model. Horizontal red lines mark the ΔAIC = 2 threshold. Numbers at the top right corner of each plot show the proportion of times the correlated model was below the ΔAIC = 2 threshold, independent of the generating model (see also Table S3).

**Table S2:** Proportion of support for each of the models across 100 simulation replicates for each of the three generating models with data sets showing the same number of species and sequence positions of the empirical data. Models were considered supported if ΔAIC < 2 units (also showed as the red horizontal line on Figure S16).

| Fitted models | Generating (true) models |  |  |
| --- | --- | --- | --- |
|  | Homogeneous | Gamma | Correlated |
| Homogeneous | 0.95 | 0.97 | 0.86 |
| Gamma | 0.00 | 0.00 | 0.00 |
| Correlated | 0.06 | 0.07 | 0.22 |

### 3.8) Model-averaged transition rates estimated across sequence positions

Figures below show the distribution of transition rates estimated along the sequence positions of the alignments. These are complementary to Figure 2 and 3 of the main text. Figures 2 and 3 show only one single alignment as a representative of the results whereas here we summarize the variance of model-averaged transition rates estimated along sequence positions. The results shown below include phylogenetic uncertainty (Figure S17), alignment uncertainty (Figure S18), as well as variation of song bout (Figure S19 and S20).

**Figure S16:**
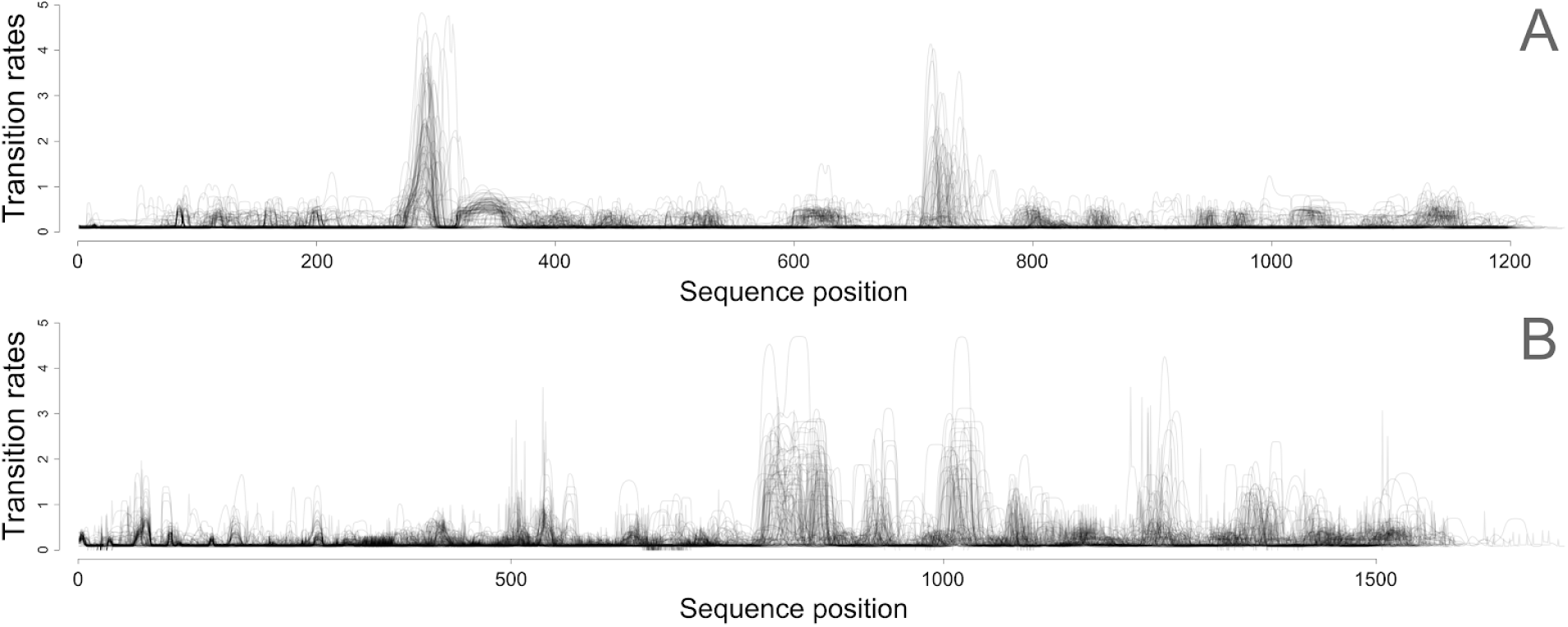
Model averaged rates of evolution estimated for the complete song sequences of *Gryllus* species under the correlated model using MAFFT conditioned on a random sample of 100 trees. A: Untransformed sequences. B: Rescaled sequences.

**Figure S17:**
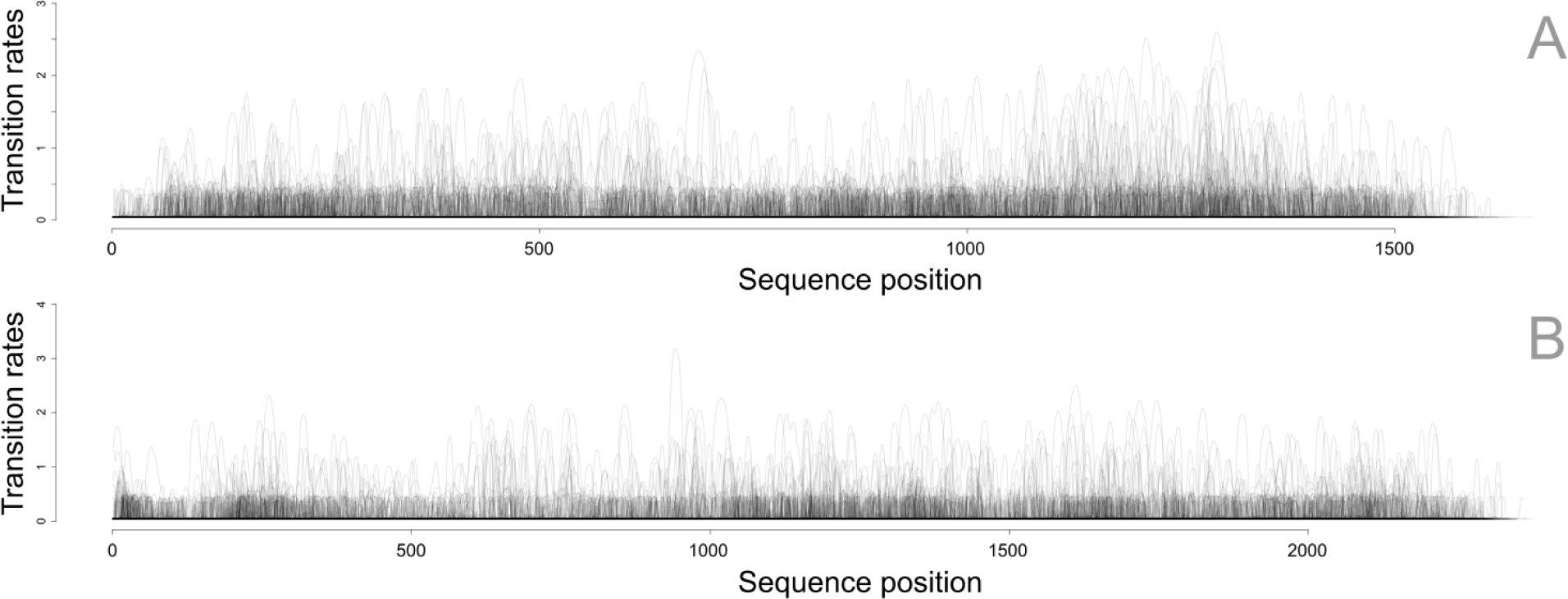
Model averaged rates of evolution estimated for the complete song sequences of *Gryllus* species using BAli-Phy conditioned on the MCC tree. Plots show random samples of 100 alignments from the posterior distribution. A: Untransformed sequences B: Rescaled sequences.

**Figure S18:**
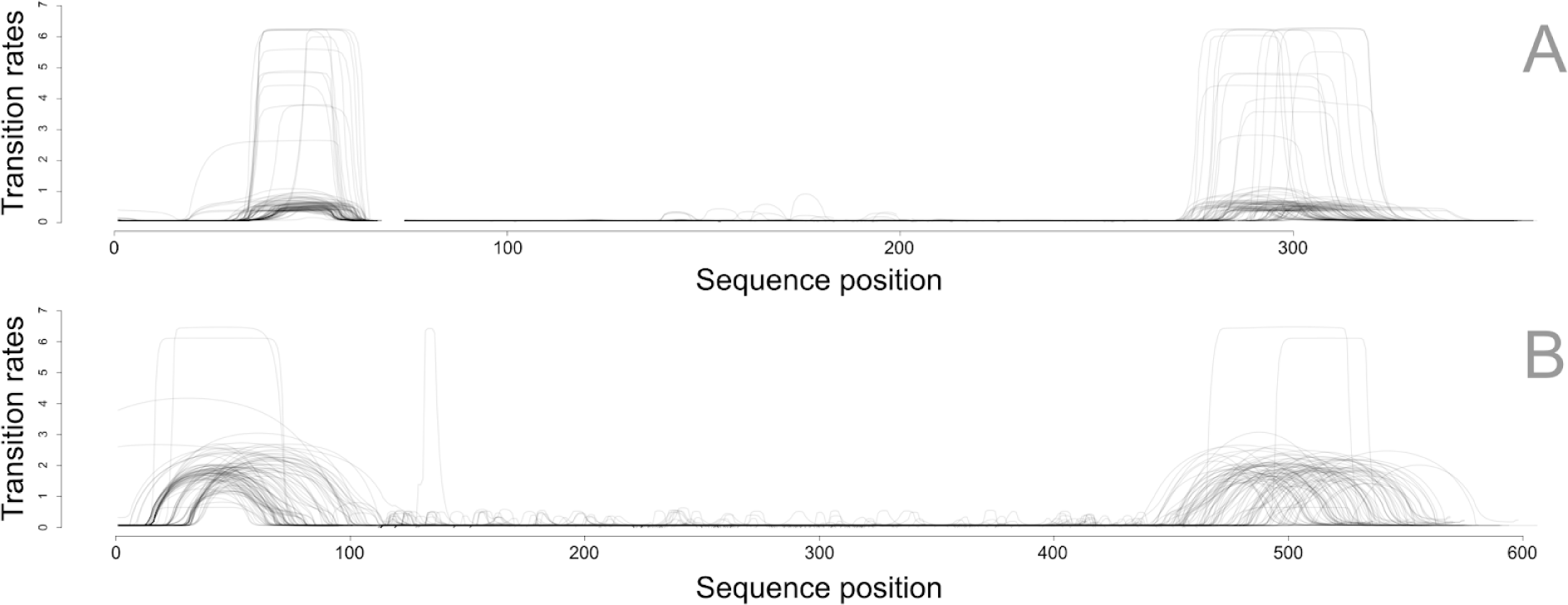
Model averaged rates of evolution estimated from a sample of 100 song bout data sets representing within species variation of songs. Alignments were estimated using MAFFT conditioned on the MCC tree. A) Original bout sequences (DO2). B) Time-rescaled bout sequences.

**Figure S19:**
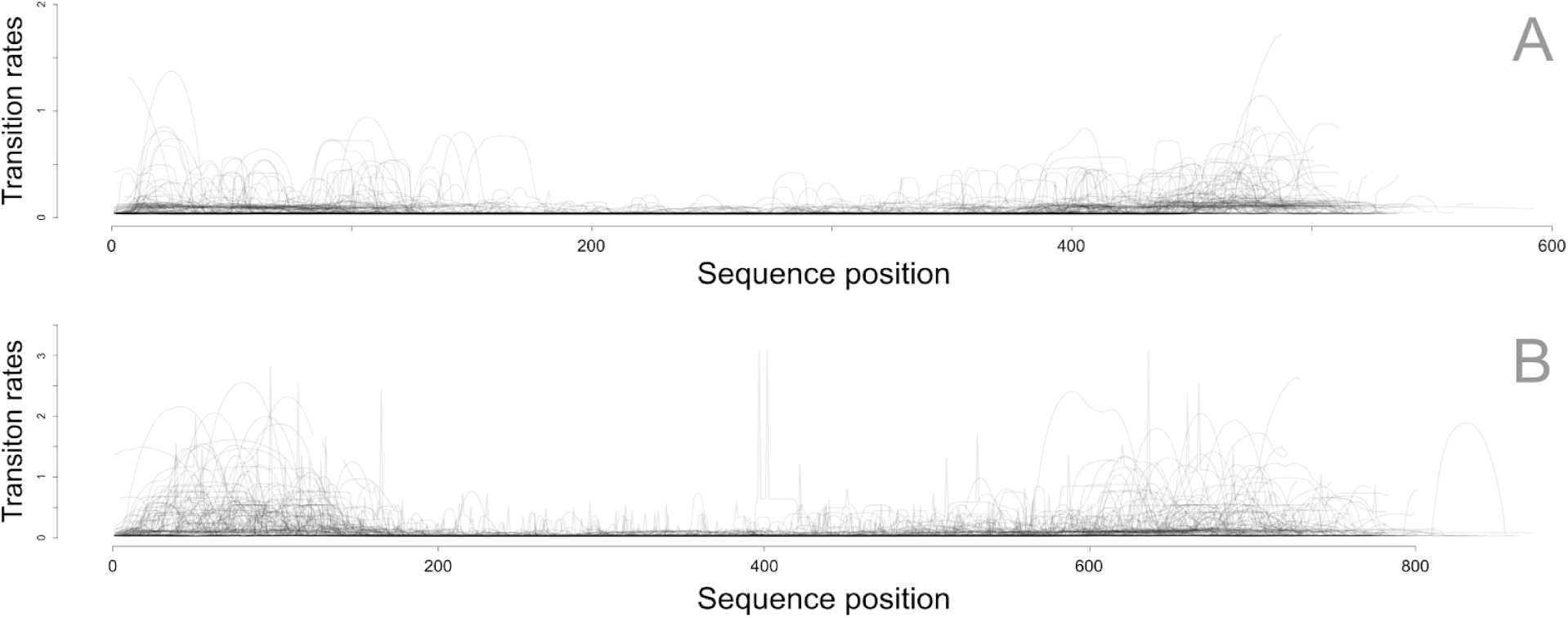
Rates of evolution estimated for the song sequence with a single song bout per species. Different from Figure S18, here only a single song bout data was used and the distribution of estimates reflects alignment uncertainty described by the posterior distribution from BAli-Phy. A) Original bout sequences (DO2). B) Time-rescaled bout sequences.

### 2.9) Comparing model-averaged rates between chirp and interchirp silence regions

In order to compare rates of transition estimated for chirp regions and interchirp silence regions we selected and grouped positions of the alignment between the groups. We selected a given alignment position as belonging to a chirp region (i.e., the column of the alignment) if more than 50% of the lineages (i.e., rows of the alignment) showed a sound character state. Likewise, we selected silence regions as positions with more than 50% silence character states. Then we recorded the distribution of model-averaged log transition rates for the positions of each group and computed median rate for each group. Here we used the median because the distribution of rates does not follow a Gaussian distribution. We repeated this operation across 100 randomly sampled alignments from the posterior distribution of the BAli-Phy analyses (i.e., incorporating alignment uncertainty) using both the time rescaled results for the whole sequence and the song bouts. We compared the median for the groups across samples using a paired Wilcoxon signed-rank test.

